# Melanotransferrin Functions as a Pro-Oncogenic WNT Agonist: A Yin-Yang Relationship in Melanoma with the WNT Antagonist and Metastasis Suppressor, NDRG1

**DOI:** 10.1101/2023.02.27.530353

**Authors:** J. Paluncic, M. Gholam Azad, D.J.R Lane, J. Skoda, K.C. Park, S. Chiang, D.H. Bae, R. Scolyer, R. Afroz, G. Babu, J. Wilmott, K. Loh, P.J. Jansson, M. Dharmasivam, M.L. Huang, X. Zhao, Z. Kovacevic, D.R. Richardson

**Affiliations:** Molecular Pharmacology and Pathology Program, Department of Pathology and Bosch Institute University of Sydney, Sydney, New South Wales, 2006, Australia; Department of Translational Medical Oncology, National Center for Tumor Diseases Dresden, Dresden, Germany; Centre for Cancer Cell Biology and Drug Discovery, Griffith Institute for Drug Discovery, Griffith University, Nathan, Brisbane, Queensland, 4111, Australia; Department of Experimental Biology, Faculty of Science, Masaryk University, Brno, Czech Republic; International Clinical Research Center, St. Anne’s University Hospital Brno, Brno, Czech Republic; Melanoma Institute Australia, Sydney Medical School, The University of Sydney, Sydney, Australia; Department of Tissue Pathology and Diagnostic Oncology, Royal Prince Alfred Hospital and New South Wales Health Pathology, Sydney, Australia; Department of Physiology, School of Biomedical Sciences, University of NSW, NSW 2052 Australia; Department of Pathology and Biological Responses, Nagoya University Graduate School of Medicine, Nagoya 466-8550, Japan.

## Abstract

A persistent mystery in the melanoma field has been the function of one of the first melanoma tumor antigens characterized, namely p97 (melanotransferrin; MTf). While MTf expression increases melanoma cell proliferation, migration, and tumorigenesis, the molecular mechanism responsible is unknown. On the other hand, N-myc down-stream regulated gene 1 (NDRG1) is a potent metastasis suppressor and WNT antagonist. Expression of NDRG1 in melanoma cells suggests a role in inhibiting metastasis, with this study investigating MTf’s role in oncogenic signaling. We demonstrate MTf acts as a pro-oncogenic WNT agonist, which down-regulates NDRG1, while silencing *MTf* increases NDRG1 expression. In contrast, silencing *NDRG1* increases MTf expression. These observations demonstrate a bidirectional negative feedback loop and “Yin-Yang” relationship between MTf and NDRG1. Mechanistically, MTf was directly associated with the WNT co-receptor, lipoprotein-receptor 6 (LRP6), and increased total LRP6 expression, activated p-LRP6 (Ser1490), β-catenin, and activated β-catenin (Ser552) levels, with MTf expression inducing their nuclear accumulation. Additionally, MTf expression increased downstream WNT targets, namely cyclin D1 and c-Myc, with c-Myc down-regulating NDRG1 expression. Silencing *c-Myc* prevented the Yin-Yang relationship between NDRG1 and MTf, indicating c-Myc played a key role in their inverse regulation. Melanoma patient specimens demonstrated that a low NDRG1/MTf ratio was significantly (*p* = 0.008) associated with lower survival and metastasis. Chemotherapeutic agents that up-regulated NDRG1 depressed MTf and nuclear LRP6 and potently inhibited melanoma xenograft growth *in vivo*. This study demonstrates MTf acts as a WNT agonist, with a Yin-Yang relationship being observed with the WNT antagonist, NDRG1.

## Introduction

Melanoma is one of the most aggressive and treatment-resistant human cancers.^1^ Melanotransferrin (MTf; Uniprot P08582; also known as p97, MFI2, and CD228) was one of the first well-characterized oncofetal antigens in melanoma.^1–3^ The highest MTf expression was demonstrated in melanoma cells, with small quantities in normal tissues.^1, 4–7^ The protein is a transferrin homolog that binds one iron atom and is mainly membrane bound *via* a glycosylphosphatidylinositol (GPI) anchor.^1, 8^ While being a transferrin homolog, recent structural studies demonstrate a novel interdomain arrangement *versus* other transferrin family members, suggesting a different function.^9^

Comprehensive studies by our laboratory using melanoma cells *in vitro*^6, 10^ and knockout and transgenic mice *in vivo*,^11–13^ demonstrated MTf played no significant role in iron homeostasis. However, silencing *MTf* in melanoma cells decreased proliferation, migration, DNA synthesis, and melanoma xenograft initiation and growth *in vivo*.^12, 14^ Across five expression models, transcription factor 4 (TCF4) was identified to be modulated by MTf.^12, 14^ This is significant, as TCF4 interacts with β-catenin to transcribe oncogenes involved in WNT signaling and proliferation.^15, 16^ Further, MTf was demonstrated to interact with an unidentified member of the lipoprotein receptor-related protein (LRP) family that play key roles in cell signaling and trafficking, with this interaction being followed by MTf internalization.^17, 18^ However, the exact molecular function and role of MTf remains unclear. Key to melanoma progression, phenotypic switching occurs between proliferation and invasion, with this being mediated by WNT signaling and alterations in TCF4 and the β-catenin-binding transcription factor, lymphoid enhancer-binding factor-1 LEF1.^19^ Considering the marked regulation of TCF4 by MTf,^14^ it was hypothesized that MTf modulates WNT signaling.

N-myc downstream regulated gene 1 (NDRG1) is a potent metastasis suppressor involved in inhibiting spread of prostate, pancreatic cancer, colon cancer, *etc*.,^20–22^ although a role in melanoma has yet to be intensively investigated. NDRG1 effectively suppresses a range of oncogenic signaling pathways, including the WNT pathway.^23–30^ In fact, NDRG1 acts as a WNT antagonist and inhibits the epithelial mesenchymal transition (EMT), by increasing plasma membrane β-catenin and preventing its nuclear translocation.^23, 27^

Herein, we demonstrate using melanoma cells that MTf expression acts as a pro-oncogenic WNT agonist that inhibits the expression of the WNT antagonist, NDRG1. In fact, an inverse “Yin-Yang” relationship between MTf and NDRG1 expression is identified that is regulated by c-Myc. MTf associates with LRP6 and β-catenin, leading to their nuclear accumulation and the activation of WNT signaling. Innovative NDRG1-inducing agents suppress MTf expression and melanoma xenograft growth with the MTf-NDRG1 Yin-Yang relationship also being identified in melanoma patient specimens. In fact, a low NDRG1/MTf ratio was significantly associated with lower survival and metastatic disease. Knowledge of the NDRG1-MTf Yin-Yang relationship will facilitate the development of bespoke melanoma therapies and prognostic markers.

## Results

### A “Yin-Yang” Relationship Between MTf and NDRG1

Considering the potential opposing roles of pro-proliferative MTf^12, 14^ and anti-oncogenic NDRG1^20–28, 30^ on WNT signaling in melanoma biology, we examined the effect of modulating MTf using multiple melanoma cell-types transfected with overexpression or silencing vectors (**Figure 1a-g**). Two different MTf overexpressing clones of SK-MEL-28 melanoma cells, namely hIF and hIE, were utilized and compared to their vector control transfected counterparts (VA). The hIF and hIE clones expressed significantly higher MTf at 90-kDa when compared to VA cells and demonstrated significantly decreased NDRG1 at 46-kDa, but not 41-kDa (**Figure 1a**). The two NDRG1 bands represent NDRG1 isoforms.^31, 32^ The upper NDRG1 band is the key, active isoform for metastasis suppression, as: **(1)** it is potently up-regulated by the specifically designed thiosemicarbazones, di-2-pyridylketone-4,4-dimethyl-3-thiosemicarbazone (Dp44mT) and di-2-pyridylketone-4-cyclohexyl-4-methyl-3-thiosemicarbazone (DpC) that inhibit tumor growth and metastasis;^30, 32, 33^ and **(2)** silencing the NDRG1 top band prevents it acting on downstream effectors (*e.g.*, E-cadherin and β-catenin) inhibiting the metastasis-inducing EMT.^23^

**Figure 1.**
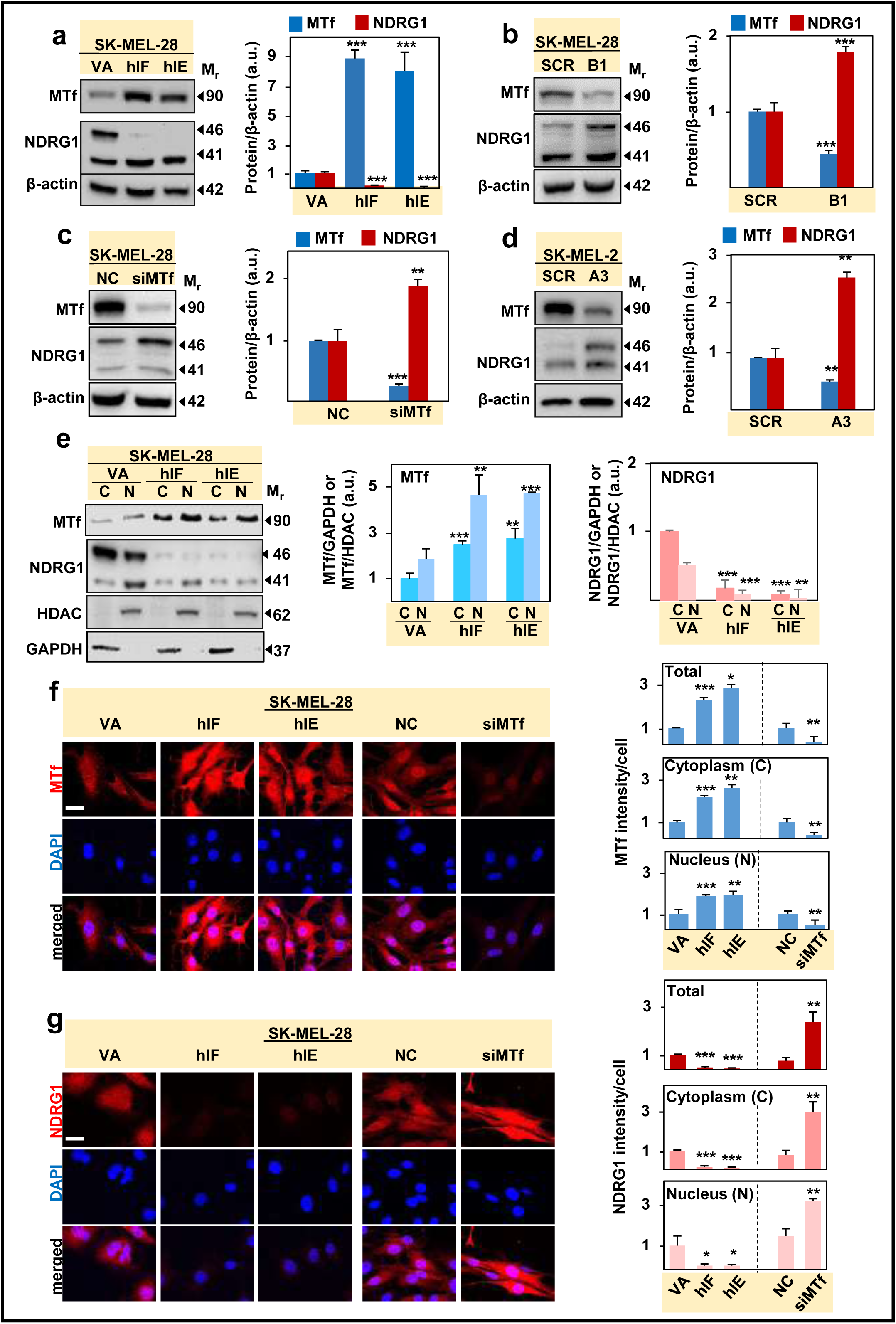
MTf overexpression (hIF and hIE clones) in SK-MEL-28 melanoma cells down-regulates NDRG1, while *MTf* silencing using SK-MEL-28 and SK-MEL-2 melanoma cells leads to NDRG1 up-regulation. Western blot analysis of: **(a)** MTf and NDRG1 expression in VA control and hIF and hIE clones; **(b)** MTf and NDRG1 expression in a stable siMTf clone (B1) of SK-MEL-28 cells *versus* the scrambled (SCR) control; **(c)** MTf and NDRG1 expression after transient transfection with siMTf in SK-MEL-28 cells *versus* the negative control (NC); **(d)** MTf and NDRG1 expression in a stable siMTf clone (A3) of SK-MEL-2 cells *versus* the SCR control; **(e)** Nuclear (N) and cytoplasmic (C) fractions of hIF and hIE clones *versus* the VA control examining MTf and NDRG1 relative to histone deacetylase (HDAC) and glyceraldehyde-3-phosphate dehydrogenase (GAPDH), respectively. **(f, g)** Confocal immunofluorescence microscopy of the VA, hIF hIE clones, as well as the NC and siMTf models examining: **(f)** MTf and **(g)** NDRG1 expression. ImageJ analysis utilized 14-31 cells over 3 experiments. Scale bar = 30 μm. Results are mean ± SD (3 experiments). \**p*<0.05; \*\**p*<0.01; \*\*\**p*<0.001 relative to control.

Similar inverse Yin-Yang regulation of MTf and NDRG1 was observed using a stable *MTf* silencing clone of SK-MEL-28 cells (*i.e.*, B1) *versus* the scrambled (SCR) control (**Figure 1b**). An additional stable *MTf* silencing clone (*i.e*., B2) ^12^ also resulted in NDRG1 up-regulation (**Supplemental Figure 1a**). This NDRG1 up-regulation was also demonstrated by transient *MTf* silencing of SK-MEL-28 cells *versus* the negative control (NC; **Figure 1c**). Stable *MTf* silencing in another melanoma cell-type, SK-MEL-2 (clone A3), also resulted in significant increase in NDRG1 (**Figure 1d**). A comparable inverse relationship between MTf and NDRG1 was also observed in 2 patient-derived melanoma cell lines (**Supplemental Figure 1b**). Collectively, these studies demonstrate that MTf has a negative effect on the metastasis suppressor, NDRG1, which may play a role in enhancing MTf’s pro-oncogenic activity.^12, 14^

### MTf and NDRG1 are Localized in the Nucleus and Cytoplasm

MTf is detected in melanoma cells as a major single band at 90-kDa (**Figure 1a-e**), with the protein being classically bound to the plasma membrane,^3^ but nuclear MTf expression has not been described. In contrast, NDRG1 demonstrates nuclear and cytosolic localization.^27, 32, 34–36^ Considering the inverse relationship between MTf and NDRG1 in multiple melanoma models (**Figure 1a-e; Supplemental Figure 1a,b)**, further studies examined the intracellular localization of NDRG1 and MTf, as this may play a role in their functions. The MTf overexpressing clones (hIF, hIE) and VA control were fractionated into their cytoplasmic (C) and nuclear (N) fractions that were validated by cytoplasmic glyceraldehyde-3-phosphate dehydrogenase (GAPDH) and nuclear histone deacetylase (HDAC) expression (**Figure 1e**). Surprisingly, MTf expression in all cell-types was almost 2-fold greater in the nuclear than cytoplasmic fractions, with MTf being significantly higher in the nuclear and cytoplasmic fractions of the hIF and hIE overexpression clones *versus* the VA control (**Figure 1e**). In contrast, total NDRG1 levels in the MTf overexpression clones was significantly lower in the nuclear and cytoplasmic fractions *versus* those in VA cells (**Figure 1e**). This localization of MTf and NDRG1 identified by cellular fractionation was verified by confocal immunofluorescence microscopy using SK-MEL-28 MTf overexpressing (hIF, hIE) and siMTf-treated cells (**Figures 1f, g**). In summary, **Figure 1** indicates a Yin-Yang relationship between NDRG1 and MTf expression.

### An Inverse Relationship Exists Between MTf Expression and NDRG1 Phosphorylation

Phosphorylation of NDRG1 (p-NDRG1) at Ser330 and Thr346 was demonstrated necessary for NDRG1 to inhibit downstream oncogenic signaling in pancreatic tumor cells.^37^ Further, the levels of these latter phosphorylations are potently increased by the agents Dp44mT and DpC that show marked anti-tumor and anti-metastatic activity.^32, 33^ As such, the effect of MTf overexpression or silencing on pNDRG1 levels was examined and again demonstrated the MTf-NDRG1 Yin-Yang relationship (**Supplemental Figures 2a-e**), which could be important for MTf’s oncogenic activity.

### *NDRG1* Silencing and Overexpression Down- and Up-Regulate MTf, Respectively

To further examine the Yin-Yang relationship between MTf and NDRG1, SK-MEL-28 (**Figure 2a-h**) or SK-MEL-2 melanoma cells (**Figure 2i**) were transfected with *NDRG1* siRNA or NC siRNA. Silencing *NDRG1* in SK-MEL-28 cells significantly decreased the 46-kDa NDRG1 band and significantly increased MTf *versus* the NC siRNA (**Figure 2a, b**). In contrast, the 41-kDa NDRG1 isoform that was reported to be consistent with a cleavage product,^31, 38^ was not silenced, as shown in other cell-types.^24, 32, 38^ Similar results were obtained for another NDRG1 siRNA (*i.e.*, siNDRG1 II; **Supplemental Figure 3a**).

**Figure 2.**
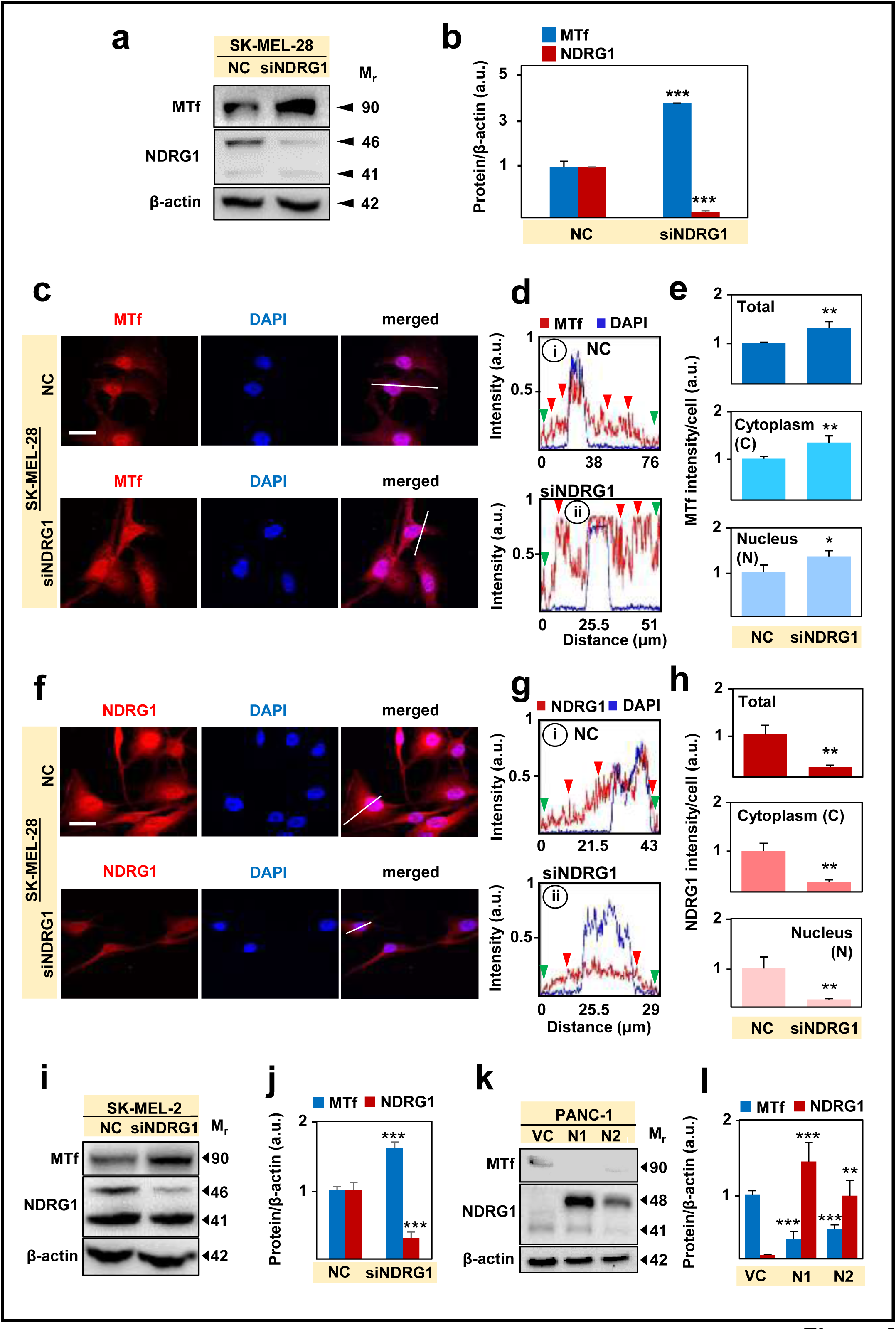
Silencing *NDRG1* in SK-MEL-28 and SK-MEL-2 melanoma cells up-regulates MTf, while NDRG1 overexpression in PANC-1 pancreatic cancer cell clones (N1 and N2) down-regulates MTf. **(a, b)** Western blot examining transient *NDRG1* silencing (siNDRG1) *versus* the negative control (NC) on MTf and NDRG1 expression in SK-MEL-28 cells. **(c-h)** Confocal immunofluorescence microscopy of SK-MEL-28 cells examining **(c-e)** MTf or **(f-h)** NDRG1 intensity and localization in cells after siNDRG1 *versus* the NC using ImageJ and its analytical tools, plot profile analysis **(di, ii; gi, ii)** and co-masking analysis **(e; h)**. Scale bar = 30 μm. The white line that crosses the cell body **(c, f)** results in the intensities of the different channels displayed in the plot profile analysis in d, g. ImageJ analysis utilized 18 cells over 3 experiments. **(i, j)** siNDRG1 increases MTf and decreases NDRG1 in SK-MEL-2 cells *versus* the NC. **(k, l)** NDRG1 overexpression decreases MTf levels in PANC-1 N1 and N2 clones *versus* vector control (VC) cells. Results are mean ± SD (3 experiments). \**p*<0.05, \*\**p*<0.01, \*\*\**p*<0.001 *versus* the respective control.

Next, confocal microscopy was performed after silencing *NDRG1* in SK-MEL-28 cells to examine MTf and NDRG1 localization (**Figure 2c-h**). Plot profile analysis of the NC **(Figure 2d(i))** demonstrated cytoplasmic (red arrows) and particularly nuclear (asterisk) MTf, with some plasma membrane MTf (green arrows). In contrast, siNDRG1 significantly increased cytoplasmic (red arrow) and nuclear MTf (asterisk; **Figure 2d(ii), e**), while decreasing total, cytoplasmic, and nuclear NDRG1 (**Fig. 2f, g(i,ii), h**). Silencing *NDRG1* in SK-MEL-2 cells also significantly up-regulated MTf and decreased NDRG1 (**Figure 2i, j**). Additional studies using pancreatic cancer cell (PANC-1) clones stably overexpressing NDRG1 (N1, N2) also significantly decreased MTf (**Figure 2k, l**), with a similar effect being demonstrated by immunofluorescence (**Supplemental Figure 3bi, ii**). Collectively, **Figures 1, 2**, and **Supplemental Figures 1-3** demonstrate an inverse Yin-Yang relationship between NDRG1 and MTf.

### MTf Overexpression or *NDRG1* Silencing Up-Regulates LRP6 and its Activation

Like other GPI-anchored proteins,^39^ MTf may function as a ligand or modulator of other transmembrane signaling molecules. Our laboratory demonstrated that in multiple MTf expression models *in vitro* and *in vivo* that this protein modulates TCF4 expression that is involved in WNT signaling.^12, 14^ Further, NDRG1 acts as a WNT antagonist by: **(1)** increasing plasma membrane β-catenin as part of the adherens complex;^23^ **(2)** inhibiting β-catenin nuclear translocation; and **(3)** directly binding the key WNT upstream co-receptor, LRP6.^40, 41^ Intriguingly, MTf has been reported to interact with an unidentified member of the LRP family.^18^ Considering this evidence and the Yin-Yang relationship between MTf and NDRG1, and the potential of both to impact WNT signaling, studies then investigated if MTf modulates the critical upstream WNT signaling protein, LRP6.

Both LRP6 and its activating phosphorylation at Ser1490, which recruits axin to the membrane to promote β-catenin signaling,^42–44^ were significantly increased upon MTf overexpression, with NDRG1 being down-regulated (**Figure 3a**). In contrast, stable (**Figure 3b**) or transient (**Figure 3c**) *MTf* silencing in SK-MEL-28 cells significantly decreased LRP6 and p-LRP6 (Ser1490) and increased NDRG1. Considering the inverse relationship between MTf and NDRG1 (**Figures 1, 2**; **Supplemental Figures 1-3**), the effects of *NDRG1* silencing and overexpression on upstream WNT signaling was then assessed (**Figure 3d; Supplemental Figure 4)**. Incubation of SK-MEL-28 cells with siNDRG1 significantly down-regulated NDRG1 and significantly increased MTf, LRP6, and p-LRP6 levels (Ser1490; **Figure 3d**). In contrast, transfection of SK-MEL-28 cells with increasing levels of an NDRG1 expression plasmid resulted in significant down-regulation of LRP6 and MTf *versus* the control (**Supplemental Figure 4a**). To further understand the mechanism involved in the up-regulation of LRP6 after MTf expression, SK-MEL-28 melanoma cells with MTf hyperexpression and the vector control VA were incubated in the presence and absence of the protein synthesis inhibitor, cycloheximide (**Supplemental Figure 4b**). These studies demonstrated the half-life of LRP6 in VA cells was significantly (*p* < 0.002) increased from 8.82 ± 0.65 h (3) to 11.50 ± 0.46 h (3) after MTf hyperexpression. Collectively, **Figure 3a-d** **and Supplemental Figure 4a, b** indicates MTf promotes upstream WNT signaling *via* increasing LRP6 levels and activation, with this being antagonized by NDRG1, elucidating a potential mechanism how MTf promotes tumorigenesis.

**Figure 3.**
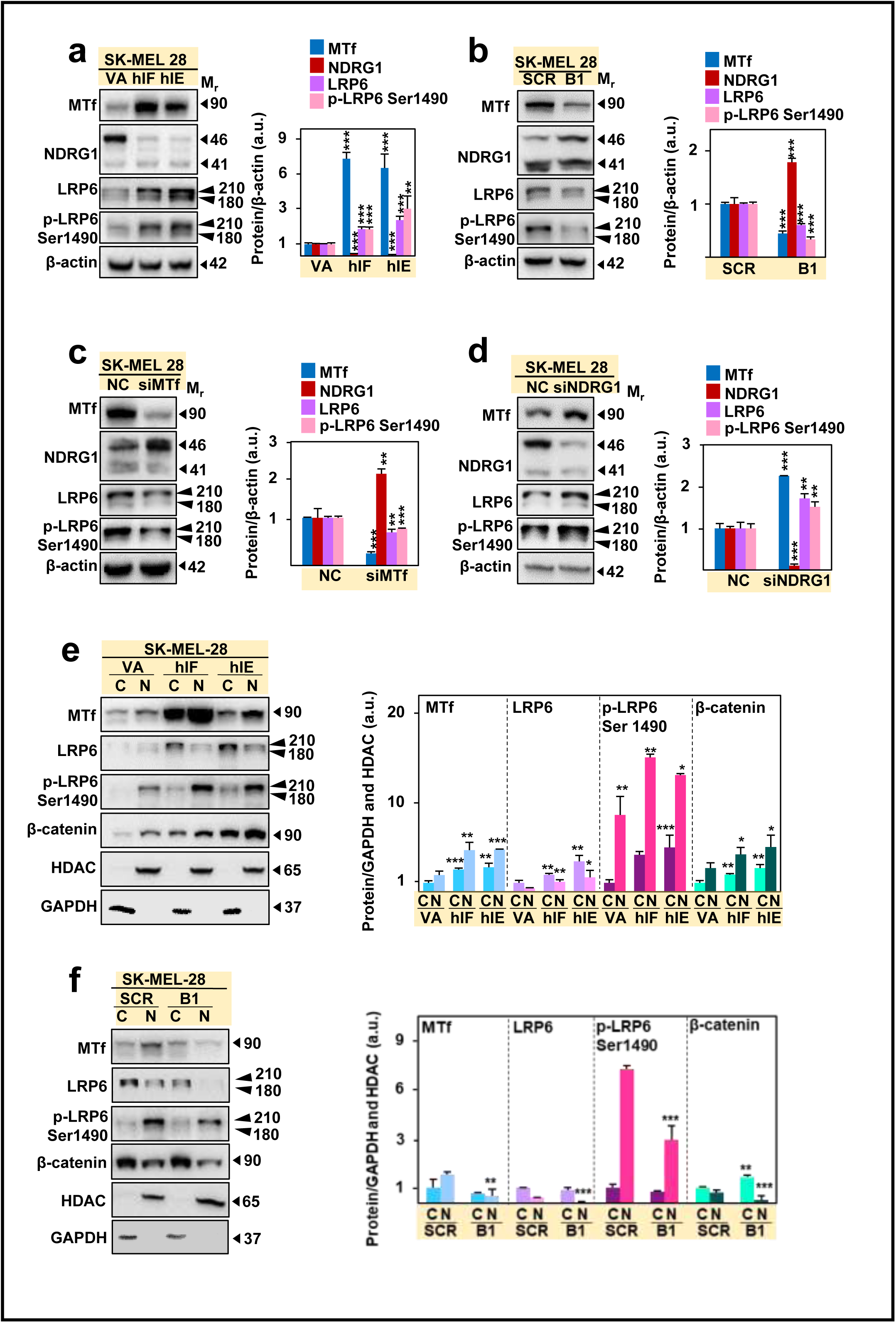
Overexpression of MTf in SK-MEL-28 melanoma cells up-regulates the expression and the nuclear translocation of LRP6, p-LRP6 (Ser1490), and β-catenin, while silencing *MTf* has an opposite effect. Western blot analysis of: **(a)** MTf, NDRG1, LRP6 and p-LRP6 (Ser1490) levels in VA control and MTf overexpressing, hIF and hIE SK-MEL-28 melanoma cell clones; **(b)** MTf, NDRG1, LRP6 and p-LRP6 (Ser1490) levels in the stable siMTf clone (B1) of SK-MEL-28 cells *versus* the scrambled (SCR) control; **(c)** MTf, NDRG1, LRP6 and p-LRP6 (Ser1490) levels after transient transfection with siMTf in SK-MEL-28 cells *versus* the negative control (NC); **(d)** MTf, NDRG1, LRP6 and p-LRP6 (Ser1490) levels after silencing *NDRG1* in SK-MEL-28 melanoma cells. **(e-f)** Western blot analysis of nuclear (N) and cytoplasmic (C) fractions of: **(e)** the MTf overexpressing, hIF and hIE clones, *versus* the VA control; and **(f)** the stable *MTf* silenced B1 clones *versus* the SCR control, examining MTf, LRP6, p-LRP6 (Ser1490) and β-catenin in these fractions *versus* the nuclear and cytoplasmic markers, histone deacetylase (HDAC) and glyceraldehyde-3-phosphate dehydrogenase (GAPDH), respectively. Results are mean ± SD (3 experiments). \**p*<0.05 \*\**p*<0.01 \*\*\**p*<0.001 *versus* the control.

### LRP6, p-LRP6 (Ser1490) and their Downstream Target, β-Catenin, are Localized in the Nucleus and Cytoplasm upon MTf Expression

Since MTf was surprisingly identified in the nucleus (**Figure 1e, f**) and increases total LRP6 and its active form, p-LRP6 (Ser1490; **Figure 3a-f**), studies further investigated if MTf regulates their nuclear translocation. Of note, LRP6 has been identified in the nucleus of normal and neoplastic cell-types where multiple effector functions have been reported.^45–47^ The current studies demonstrate LRP6 and p-LRP6 (Ser1490) were significantly increased in the cytoplasmic (C) and nuclear (N) fractions of the MTf overexpression clones (**Figure 3e**). MTf overexpression also significantly increased nuclear β-catenin translocation and its cytoplasmic levels (**Figure 3e**). In contrast, stable silencing of *MTf* in the B1 clone of SK-MEL-28 cells significantly decreased MTf, LRP6, and p-LRP6 (Ser1490) in the nuclear fraction *versus* the nuclear SCR control (**Figure 3f**). Stable *MTf* silencing also significantly decreased nuclear β-catenin *versus* the SCR control, while the cytoplasmic fraction was significantly increased compared to the SCR control cytoplasmic fraction (**Figure 3f**).

In conclusion, **Figure 3e,f** demonstrates that MTf overexpression increases nuclear translocation of LRP6, p-LRP6 (Ser1490), and β-catenin, while silencing *MTf* reverses this effect. Considering the role of activated p-LRP6 (Ser1490) in WNT signaling,^48^ these data suggest MTf enhances WNT signaling.

### MTf Associates with LRP6, p-LRP6 (Ser1490), β-catenin, and p-β-catenin (Ser552)

Given MTf’s role in up-regulating total and activated levels of LRP6, β-catenin, and their nuclear translocation, it was crucial to mechanistically dissect how MTf regulates these proteins. Co-immunoprecipitation of MTf using an anti-MTf antibody demonstrated a significant increase in MTf, LRP6, p-LRP6 (Ser1490), β-catenin, and p-β-catenin (Ser552) in MTf overexpressing cells *versus* VA control cells (**Figure 4a**). In contrast, stable silencing of *MTf* in B1 cells resulted in a significant decrease in the association of MTf with LRP6, p-LRP6 (Ser1490), β-catenin and p-β-catenin (Ser552) relative to the SCR control (**Figure 4b**). Together, since nuclear p-β-catenin (Ser552) and p-LRP6 (Ser1490) activate WNT signaling,^48, 49^ **Figure 4a,b** suggests an association of MTf with LRP6 and β-catenin to promote oncogenic signaling.

**Figure 4.**
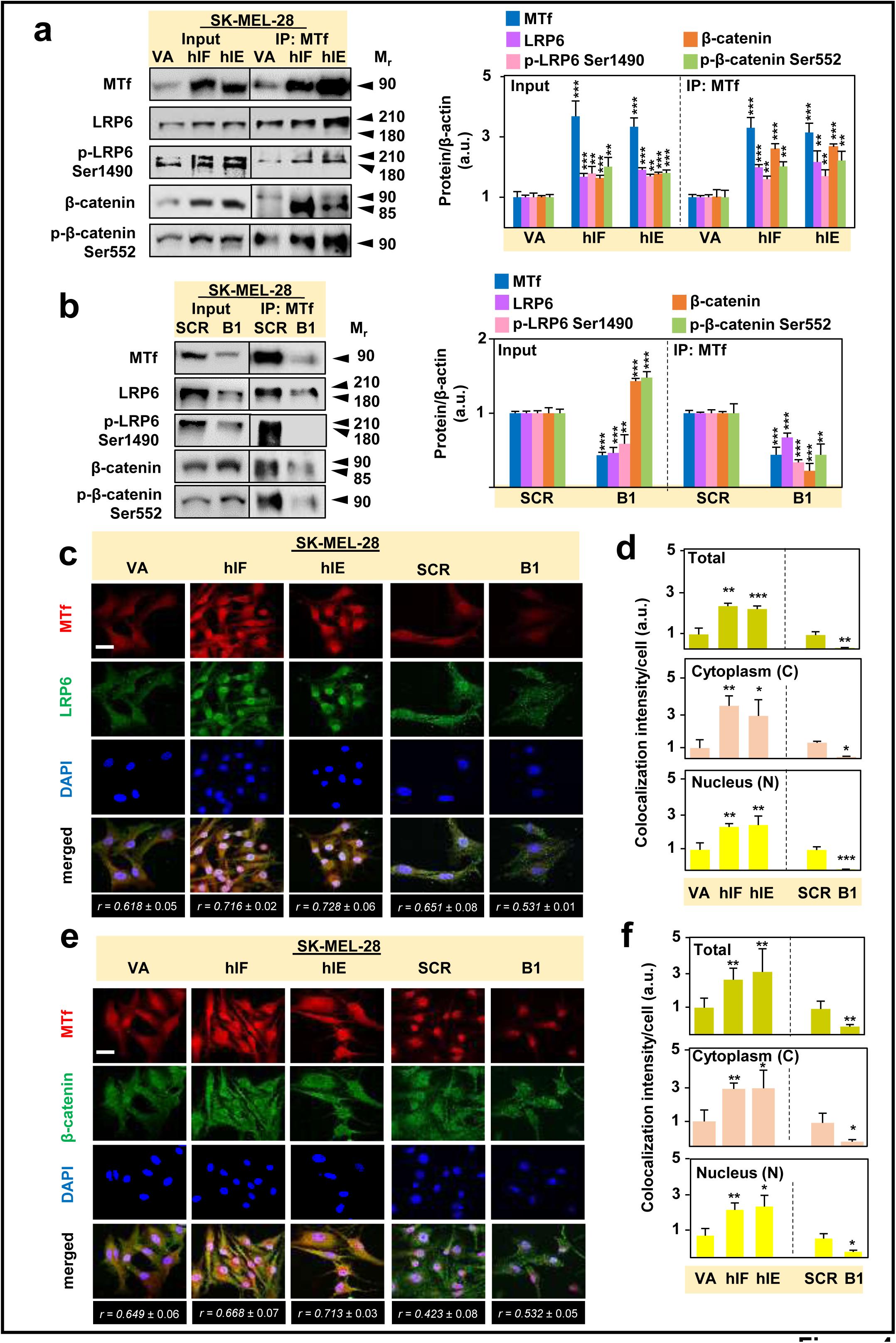
MTf associates with LRP6, p-LRP6 (Ser1490), β-catenin and p-β-catenin S552 in SK-MEL-28 melanoma cells, with MTf overexpression increasing their association and leading to increased nuclear LRP6 and β-catenin levels and their co-localization with MTf. **(a, b)** MTf was co-immunoprecipitated and western blotting performed in SK-MEL-28 cells to examine: MTf, LRP6, p-LRP6 (Ser1490), β-catenin and p-β-catenin S552 using: **(a)** MTf overexpressing hIF and hIE clones *versus* the VA control; and **(b)** the stable *MTf* silenced B1 clones *versus* the SCR control, **(c-e)** Confocal immunofluorescence microscopy was performed using hIE and hIF MTf overexpressing cells *versus* VA control cells relative to *MTf* silenced B1 cells *versus* SCR control to examine: **(c, d)** Co-localization between MTf and LRP6 and **(e, f)** co-localization between β-catenin and MTf. The ImageJ co-localization analysis tool was used in **(d)** and **(f)** and utilized 11-52 cells over 3 experiments. Scale bar = 30 μm. Results are mean ± SD (3 experiments). \**p*<0.05, \*\**p*<0.01, \*\*\**p*<0.001 relative to the control.

To additionally assess this interaction, when LRP6 was co-immunoprecipitated and assessed for LRP6 and MTf by western analysis, a significant increase of both was observed in MTf overexpressing cells (**Supplemental Figure 5a**). In contrast, examining LRP6 and MTf levels when LRP6 was immunoprecipitated from stable *MTf* silenced cells, both LRP6 and MTf were significantly decreased in the immunoprecipitate (**Supplemental Figure 5b**). When an anti-β-catenin antibody was used to immunoprecipitate β-catenin from MTf overexpressing cells, both MTf and β-catenin were at higher levels than the VA control (**Supplemental Figure 5c**). An opposite effect was demonstrated for the *MTf* silencing clone, where upon co-immunoprecipitation with the anti-β-catenin antibody, MTf was significantly lower in the immunoprecipitate (**Supplemental Figure 5d**). Collectively, these co-IP studies in **Figure 4** and **Supplemental Figure 5** demonstrate an association between MTf and the WNT upstream and downstream effectors, namely LRP6 and β-catenin, plus their activated forms, that were increased upon MTf overexpression.

### MTf Overexpression Increases LRP6 Co-Localization with MTf and its Nuclear Translocation

To further investigate MTf association with LRP6 or β-catenin and its activated forms, confocal microscopy was used. Co-localization between MTf and LRP6 was significantly increased in the over-MTf overexpression clones (hIF and hIE; **Figure 4c, d; Supplemental Figure 5e**), with co-localization also evident for MTf and β-catenin (**Figure 4e, f; Supplemental Figure 5f**). This resulted in yellow vesicular-like staining in the cytoplasm and white (LRP6/MTf) or purple (β-catenin/MTf) granular nuclear staining (**Supplemental Figure 5e, f**). Comparing co-localization of MTf and either LRP6 or β-catenin in *MTf* silenced B1 cells, there was a significant decrease in total, cytoplasmic and nuclear co-localization (**Figure 4c, d; Supplemental Figure 5e**).

Examining p-LRP6 (Ser1490) in the MTf overexpressing clones (hIF and hIE) and *MTf* silenced B1 clones using confocal microscopy (**Supplemental Figure 6a**) demonstrated MTf overexpression increased total, nuclear, and cytoplasmic p-LRP6 (Ser1490), while *MTf* silencing had the opposite effect using plot profile analysis (**Supplemental Figure 6b**) or quantification of intensity (**Supplemental Figure 6c**). In summary, **Supplemental Figure 6** demonstrated that MTf overexpression resulted in increased nuclear translocation of the activated LRP6 (p-LRP6 (Ser1490)) WNT co-receptor involved in WNT signaling.^48^

Collectively, MTf expression increased nuclear translocation of LRP6, p-LRP6 (Ser1490), and β-catenin that is involved in WNT oncogenic signaling.^48^ These data demonstrate a novel association between MTf and both LRP6 and β-catenin and their nuclear co-localization upon MTf expression.

### MTf Overexpression Up-Regulates the β-Catenin Downstream Targets, c-Myc and cyclin D1

The EMT is a critical step in metastasis,^50^ with WNT signaling playing a key role.^19, 51–53^ NDRG1 inhibits the EMT by: **(1)** stabilizing the adherens junction composed of E-cadherin and β-catenin, at the cell membrane;^23, 27^ and **(2)** preventing β-catenin oncogenic nuclear translocation,^23, 27^ that transcribes *c-Myc* and *cyclin D1*.^54–56^ As MTf down-regulates NDRG1 (**Figure 1a, e, g**), the effect of MTf overexpression on the WNT signaling pathway in melanoma cells was assessed. Expression of MTf significantly increased β-catenin and phosphorylated β-catenin (serine 552; p-β-catenin S552), which increases its transcriptional activity,^49^ and also significantly increased the WNT effectors, c-Myc and cyclin D1 (**Fig. 5a**). Confocal microscopy demonstrated that MTf overexpression significantly increased total, cytoplasmic, and nuclear cyclin D1, while siMTf decreased this (**Supplemental Figure 7a, b, c**).

**Figure 5.**
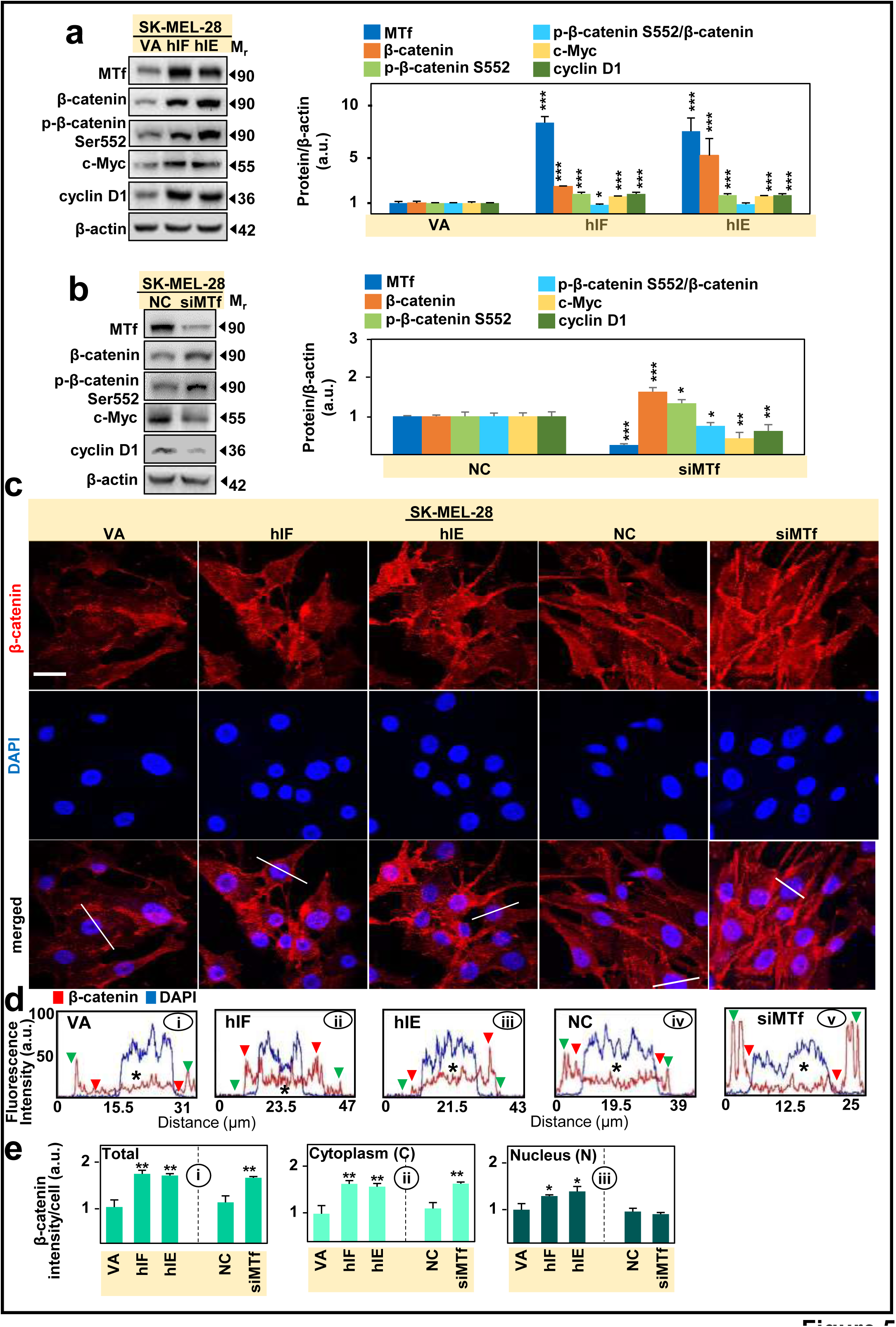
MTf overexpression in SK-MEL-28 melanoma cells increases total and nuclear β-catenin, p-β-catenin (Ser552), and the downstream effectors of WNT signaling, c-Myc and cyclin D1. **(a)** Western blotting demonstrates that MTf overexpression in SK-MEL-28 cells (hIF and hIE clones) increases total β-catenin, p-β-catenin S552, c-Myc and cyclin D1 levels *versus* VA controls. **(b)** Silencing *MTf* (siMTf) in SK-MEL-28 cells increases total β-catenin, p-β-catenin S552 and decreases c-Myc and cyclin D1 *versus* the negative control (NC). **(c-e)** Confocal immunofluorescence microscopy examining the subcellular localization of β-catenin in SK-MEL-28 MTf overexpressing hIE and hIF clones *versus* VA control cells or after siMTf treatment *versus* the NC. ImageJ plot profile analysis **(d)** and co-masking analysis **(e)** of the images in **(c)**. The white line that crosses the cell body in **(c)** results in the intensities of the different channels displayed in the plot profile. ImageJ analysis utilized 27-55 cells over 3 experiments. Scale bar = 30 μm. Results are mean ± SD (3 experiments). \**p*<0.05, \*\**p*<0.01, \*\*\**p*<0.001 relative to the respective control.

Upon transient *MTf* silencing in SK-MEL-28 cells, there was a significant increase of total β-catenin and p-β-catenin (Ser552), while c-Myc and cyclin D1 levels significantly decreased (**Figure 5b**). Similar results were obtained with the stable *MTf* silencing clones, B1 and B2, where total β-catenin was up-regulated (**Supplemental Figure 7b**), and c-Myc and cyclin D1 were down-regulated (**Supplemental Figure 7c).** As discussed above, the cellular localization of β-catenin, rather than its total levels, are crucial to investigate in terms of its oncogenic function.^23, 27, 54–56^ Confocal microscopy and plot profile analysis of the vector control VA (**Figure 5c, d(i)**) and NC cells (**Figure 5d(iv)**) demonstrated β-catenin was highly expressed on the plasma membrane (green arrows) with less cytoplasmic (red arrows) and also nuclear localization (asterisk; **Figure 5d(i, iv)**). In contrast, MTf overexpression in hIF and hIE cells increased nuclear (asterisk) and cytoplasmic (red arrow) β-catenin and decreased its membrane levels (**Figure 5d(ii, iii)**). In contrast, using *MTf* silenced cells (**Figure 5c**), β-catenin was prominently increased at the cell periphery in a position consistent with plasma membrane (green arrows; **Figure 5d(v)**) relative to the NC (**Figure 5d(iv)**), while being decreased in the nucleus (asterisk; **Figure 5d(v)**). Hence, MTf overexpression increases oncogenic nuclear β-catenin, while *MTf* silencing increases β-catenin levels in the plasma membrane, which is potentially anti-oncogenic due to its role in adherens junction formation.^23, 27, 57^

Analysis of **Figure 5c** demonstrated total, cytoplasmic, and nuclear β-catenin intensity was significantly increased upon MTf overexpression in the hIF and hIE clones (**Figure 5e(i-iii)**). In contrast, after *MTf* silencing, total and cytoplasmic β-catenin was significantly increased, while nuclear β-catenin was only slightly (*p* > 0.05) decreased (**Figure 5e(i-iii)**). Analysis of cellular localization of p-β-catenin (Ser552; **Supplemental Figure 8a**) demonstrated MTf overexpression in hIF and hIE cells markedly increased nuclear and cytoplasmic (red arrow) p-β-catenin (Ser552) (**Supplemental Figure 8b(ii, iii)**), while *MTf* silencing decreased nuclear p-β-catenin (Ser552; **Supplemental Figure 8b(v)**) *versus* the NC (**Supplemental Figure 8b(iv)**). Further analysis demonstrated total, cytoplasmic, and nuclear p-β-catenin (Ser552) was significantly increased upon MTf overexpression (**Supplemental Figure 8c(i-iii)**). In contrast, after *MTf* silencing, total and nuclear p-β-catenin (Ser552) was significantly decreased, while cytoplasmic p-β-catenin (Ser552) was significantly increased (**Supplemental Figure 8c(i-iii)**). Overall, **Figure 5** and **Supplemental Figure 8** demonstrate MTf expression results in β-catenin and p-β-catenin (Ser552) translocation to the nucleus and cytoplasm that could lead to the increased c-Myc and cyclin D1 observed, while *MTf* silencing acts oppositely.

### *NDRG1* Silencing Increases β-Catenin and its Nuclear Translocation

Considering the inverse relationship between NDRG1 and MTf and the results in **Figure 5** **and Supplemental Figure 8** demonstrating MTf’s effect on β-catenin, studies then examined if NDRG1 modulates β-catenin expression in SK-MEL-28 cells (**Figure 6**). Silencing *NDRG1* significantly increased MTf, β-catenin, p-β-catenin (Ser552), c-Myc and cyclin D1 (**Figure 6a, b**). Confocal microscopy (**Figure 6c**) demonstrated *NDRG1* silencing markedly decreased plasma membrane β-catenin (green arrows) and increased cytoplasmic (red arrows) and nuclear β-catenin (asterisk; ***cf.*** **Figure 6d(ii)**). There was significantly increased total, cytoplasmic and nuclear β-catenin after incubation with siNDRG1 (**Figure 6e**). Further, si*NDRG1* (**Figure 6f**) decreased cytoplasmic and plasma membrane p-β-catenin (Ser552), with a significant increase in the nucleus (**Figure 6g, h**). In summary, **Figure 6** demonstrates *NDRG1* silencing up-regulates MTf and increases nuclear β-catenin and p-β-catenin (Ser552), indicating promotion of WNT signaling.

**Figure 6.**
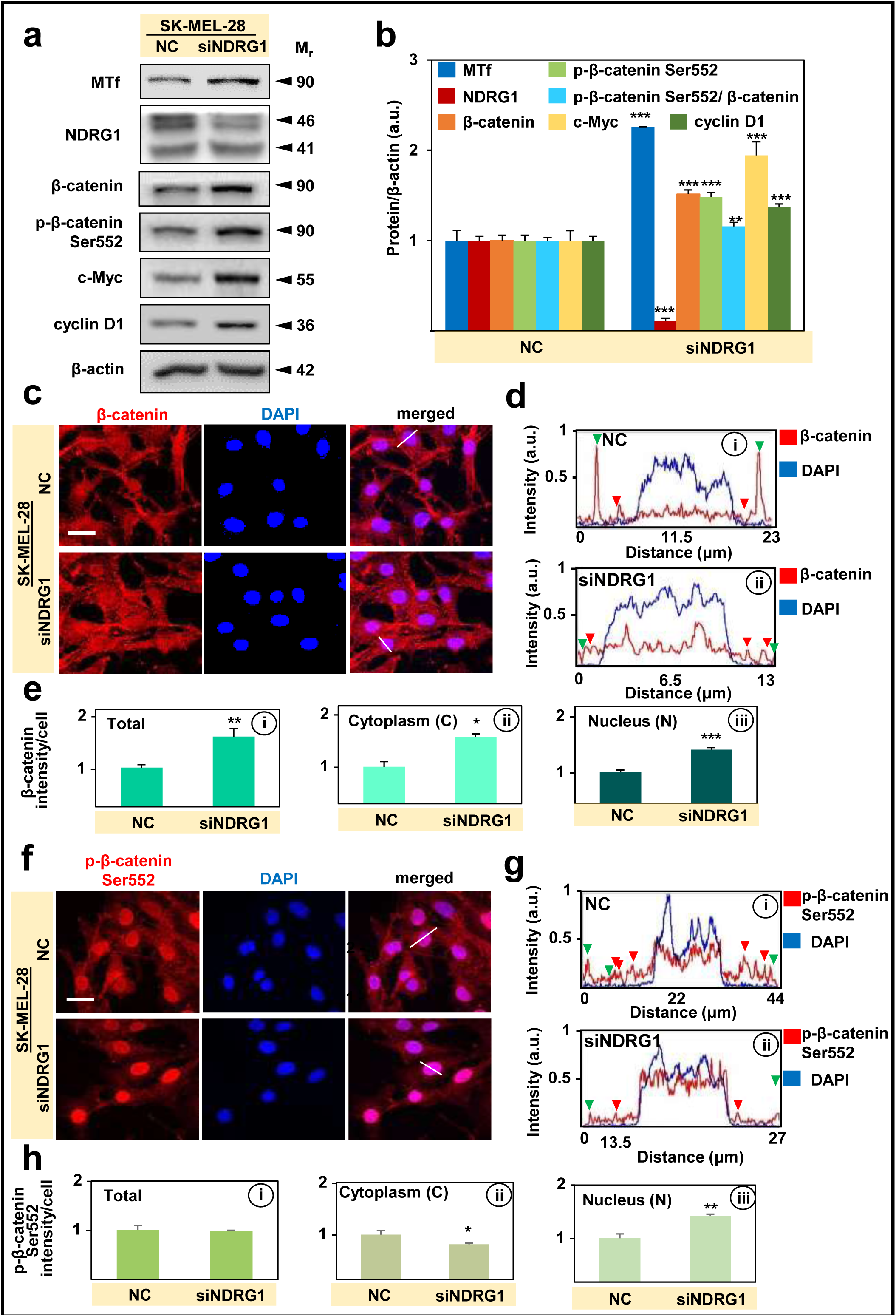
Silencing *NDRG1* in SK-MEL-28 melanoma cells results in β-catenin and p-β-catenin S552 nuclear translocation and up-regulation of c-Myc and cyclin D1. **(a, b)** Transient silencing of *NDRG1* (siNDRG1) relative to the negative control (NC), up-regulates MTf, total β-catenin, p-β-catenin S552, c-Myc and cyclin D1, as shown by western blotting. **(c-h)** Confocal immunofluorescence microscopy examining the subcellular localization of β-catenin **(c)** or p-β-catenin S552 **(f)** in SK-MEL-28 cells after treatment with siNDRG1 or the NC. ImageJ plot profile analysis **(d, g)** and co-masking analysis **(e, h)** of the images in **(c, f)**. The white line that crosses the cell body **(c, f)** results in the intensities of the different channels displayed in the plot profile. ImageJ analysis utilized 25-27 cells over 3 experiments. Scale bar = 30 μm. Results are mean ± SD (3 experiments). \**p*<0.05, \*\**p*<0.01, \*\*\**p*<0.001 relative to the respective control.

### MTf Increases *c-Myc* mRNA and Protein and its Nuclear Translocation, while Silencing *c-Myc* Negates the Ability of MTf to Down-Regulate NDRG1

Considering MTf increases WNT signaling to up-regulate c-Myc (**Figure 5a**), studies then assessed the effect of MTf overexpression and silencing on *NDRG1, LRP6,* and *c-Myc* mRNA levels (**Figure 7a**). Upon MTf overexpression, *MTf, LRP6* and *c-Myc* mRNA levels were significantly increased, while *NDRG1* mRNA was significantly decreased. The B1 *MTf* stable silencing clone demonstrated opposite effects to MTf overexpression (**Figure 7a**). Hence, MTf regulates *NDRG1*, *LRP6* and *c-Myc* at the mRNA and protein levels.

**Figure 7.**
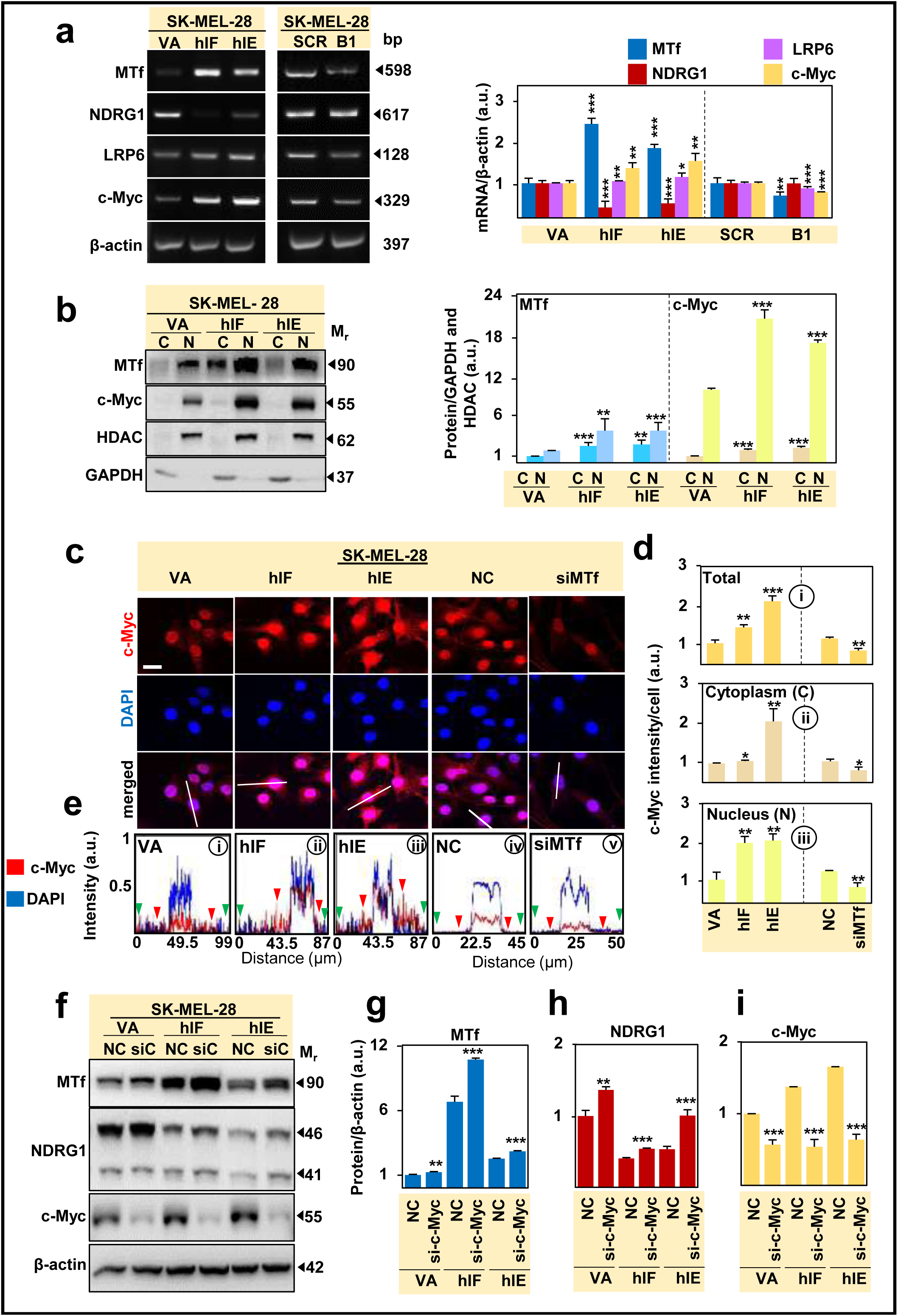
(a-e) Overexpression of MTf in SK-MEL-28 melanoma cells (hIF and hIE) up-regulates *MTf*, *NDRG1*, *LRP6* and *c-Myc* mRNA, and induces c-Myc nuclear translocation. (f-i) Silencing *c-Myc* in VA, hIF and hIE clones negates the MTf-NDRG1 Yin-Yang relationship. **(a)** The effect of MTf overexpression (hIF and hIE clones *versus* VA control) or stable *MTf* silencing (B1 *versus* scramble; SCR) in SK-MEL-28 cells on *MTf, NDRG1, LRP6 and c-Myc* mRNA levels using RT-PCR. **(b)** Western analysis of the nuclear (N) and cytoplasmic (C) fractions of the hIF and hIE clones *versus* the VA control examining MTf and c-Myc *versus* HDAC and GAPDH, respectively. **(c-e)** Confocal immunofluorescence microscopy examining the subcellular localization of c-Myc in SK-MEL-28 cells overexpressing MTf *versus* the VA control. ImageJ co-masking analysis **(d)** and plot profile analysis **(e)** of the images in **(c)**. The white line that crosses the cell body in the merge of **(c)** results in the intensities of the different channels displayed in the plot profile. ImageJ analysis utilized 15-39 cells over 3 experiments. Scale bar = 30 μm. **(f-i)** Transient silencing of *c-Myc* (si-C) *versus* the NC, up-regulates MTf and NDRG1 expression, as shown by western blotting. Results are mean ± SD (3 experiments). \**p*<0.05, \*\**p*<0.01, \*\*\**p*<0.001 relative to the respective control.

Since *NDRG1* mRNA levels were decreased in MTf overexpressing cells and as the known NDRG1 repressor, c-Myc,^58^ was up-regulated under these conditions (**Figures 5a and 7a**), the next step investigated c-Myc nuclear translocation. MTf overexpression in the hIF and hIE clones markedly and significantly increased nuclear c-Myc (**Figure 7b**), which was confirmed by confocal microscopy (**Figure 7c**), co-masking analysis (**Figure 7d(i-iii)**), and plot profile analysis (**Figure 7e(i-iii)**). In contrast, siMTf down-regulated total, cytoplasmic and nuclear c-Myc (**Figure 7d(i-iii)**, **7e(iv, v)**). To assess the mechanistic relationship between c-Myc, MTf, and NDRG1, silencing of *c-Myc* was investigated and led to significantly increased MTf and NDRG1 in both MTf overexpression clones and VA cells (**Figure 7f-i**). Hence, *c-Myc* silencing resulted in some loss of the inverse relationship between NDRG1 and MTf observed in the NC (**Figure 7**). Hence, these results indicate the inverse Yin-Yang relationship between NDRG1 and MTf observed under control conditions is partially disrupted by *c-Myc* silencing, indicating c-Myc plays a role in the inter-relationship between MTf and NDRG1.

### MTf Promotes Melanoma Proliferation, with Thiosemicarbazones Up-Regulating NDRG1 to Down-Regulate MTf and Inhibit Melanoma Xenograft Growth *In Vivo*

Our previous studies using melanoma and other cell-types indicated MTf is pro-proliferative *in vitro* and *in vivo,*^12, 14^ with this observation being confirmed by others.^34^ Herein, both MTf overexpression (hIF and hIE) significantly (*p*<0.001) enhanced proliferation after 3-4 days, while the stable *MTf* silencing clones, B1 and B2, significantly decreased proliferation after this time (**Supplemental Figure 9a, b**).

The studies above suggest down-regulating pro-oncogenic MTf could be an innovative therapeutic strategy. Considering this and the inverse relationship between NDRG1 and MTf (**Figure 1**), it is notable that NDRG1 is effectively up-regulated by the novel di-2-pyridylketone thiosemicarbazone (DpT) class of anti-cancer agents, such as Dp44mT and DpC,^33, 59, 60^ with DpC entering clinical trials.^61^ To examine the effect of pharmacologically up-regulating NDRG1 expression, SK-MEL-28 melanoma cells (VA) were incubated with media alone (Control), or media containing increasing concentrations of DpC or Dp44mT (1-, 5- and 10-μM) for 24 h/37°C. The expression of MTf, NDRG1, and cyclin D1 was then examined by western blotting (**Supplemental Figure 10a-d**). In these studies, DpC and Dp44mT significantly decreased MTf and cyclin D1 levels at 5-10 μM (**Supplemental Figure 10b, d**), and in agreement with previous studies,^33, 59, 62^ both agents potently up-regulated NDRG1 (**Supplemental Figure 10c**).

Similar results were also demonstrated by confocal microscopy after incubation for 24 h of SK-MEL-28 cells with Dp44mT (5 µM), where NDRG1 was significantly up-regulated, while MTf and LRP6 were significantly down-regulated (**Supplemental Figure 11**). Further, Dp44mT significantly increased the nuclear co-localization of NDRG1 and LRP6, and significantly decreased the nuclear co-localization of MTf and LRP6 (**Supplemental Figure 11**). These results after pharmacological up-regulation of NDRG1 by Dp44mT were similar to those after *MTf* silencing that up-regulates NDRG1 (**Figures 3b, c**)

To examine the anti-tumor activity of Dp44mT *in vivo*, SK-MEL-28 melanoma xenografts were established in nude mice and then treated *i.v.* at 0.625-0.7 mg/kg/5 days/week for 2 weeks. All Dp44mT doses markedly and significantly suppressed net melanoma tumor growth (**Supplemental Figure 12**). In conclusion, pharmacological targeting of NDRG1 by Dp44mT (**Supplemental Figure 10, 11**) has the same anti-oncogenic effect on decreasing MTf levels as genetic *NDRG1* up-regulation.

### The Yin-Yang Relationship between MTf and NDRG1 is Demonstrated in Patient Samples

Next, the impact of MTf and NDRG1 expression on clinical outcomes of melanoma patients was then evaluated. Treatment naïve patient samples from the TCGA-SKCM dataset (National Cancer Institute, Genomic Data Commons Data Portal) were analyzed by taking the median value and then dividing patients into “low” or “high” *MTf* mRNA (**Figure 8a(i)**) and *NDRG1* mRNA (**Figure 8a(ii)**). The *NDRG1*/*MTf* ratio was then calculated (**Figure 8a(iii)**). These studies demonstrated that high *MTf* mRNA and low *NDRG1* mRNA levels trend non significantly towards poor survival in melanoma patients (**Figure 8a(i, ii)**). However, a low *NDRG1*/*MTf* mRNA ratio significantly (*p* < 0.008) predicts poor melanoma patient survival **(Figure 8a(iii))**.

**Figure 8.**
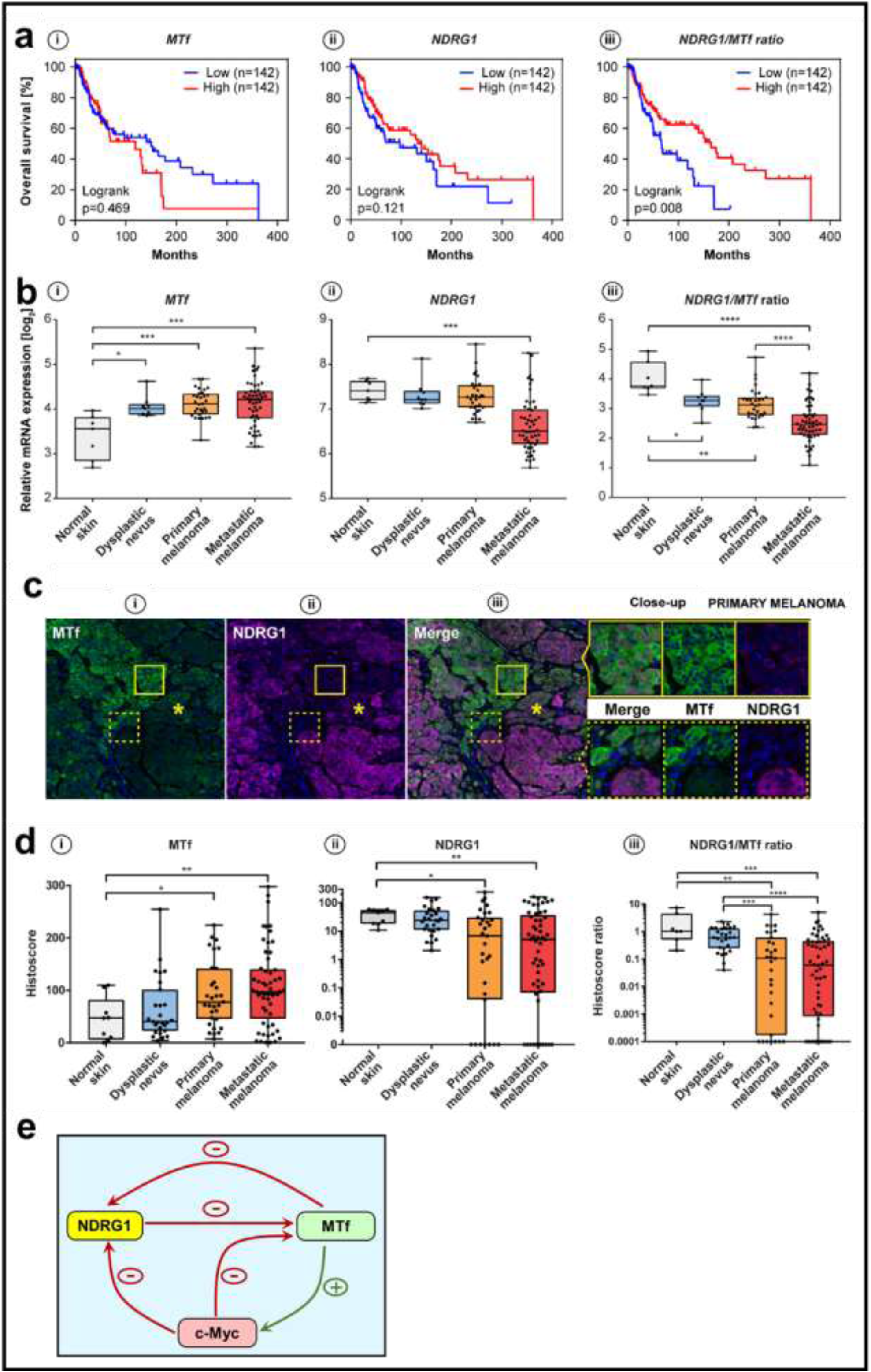
Analysis of melanoma patient tumor samples examining MTf and NDRG1 expression. **(a)** Overall survival analysis of melanoma patients based on the mRNA levels of *MTf* or *NDRG1* using treatment naïve samples from the TCGA-SKCM dataset. **(b)** Analysis of *MTf* and *NDRG1* mRNA expression in normal skin samples (*n* = 8), dysplastic naevi (*n* = 9), primary melanoma (*n* = 31), and metastatic melanoma (*n* = 52) from data provided in the GEO GSE46517 dataset. **(c)** Representative images of mIHC analysis of MTf and NDRG1 expression in primary melanoma. Melanoma tissue samples reveal a clear negative relationship between MTf and NDRG1. **(d)** MTf and NDRG1 histoscores calculated based on analysis of SOX10-positive cells in normal skin samples (*n* = 9), dysplastic naevi (*n* = 27), primary melanoma (*n* = 31) and metastatic melanoma tissues (*n* = 57). **(e)** Schematic illustrating the NDRG1-MTf Yin-Yang relationship that results from a negative bidirectional relationship between NDRG1 and MTf expression that is linked by c-myc. Results in **(b)** and **(d)** are mean ± SD. \**p*<0.05, \*\**p*<0.01, \*\*\**p*<0.001, \*\*\*\**p*<0.0001.

Considering the above, the Gene Expression Omnibus dataset (GSE46517, NCBI; **Figure 8b(i-iii)**) was then examined, which comprises normal skin (*n* = 8), dysplastic nevi (*n* = 9), and primary (*n* = 31) and metastatic melanoma (*n* = 52). Compared to normal skin, *MTf* mRNA was significantly up-regulated in dysplastic nevi and also primary and metastatic melanoma (**Figure 8b(i)**). Conversely, *NDRG1* mRNA levels were markedly decreased in metastatic melanoma (**Figure 8b(ii)**), corroborating the role of NDRG1 as a metastasis suppressor, as found in other tumor-types.^20–22^ Strikingly, *NDRG1*/*MTf* mRNA ratios were the most indicative of the progression towards the metastatic state (**Figure 8b(iii)**). While higher *NDRG1* mRNA levels were associated with low *MTf* mRNA levels in normal skin, this ratio was reversed in metastatic melanoma, which exhibited the highest *MTf* mRNA levels and lowest *NDRG1* expression. Calculation of the *NDRG1*/*MTf* mRNA ratios demonstrated there was a significant decrease between normal skin and both dysplastic nevi and primary melanoma and also a significant decrease between primary and metastatic melanoma (**Figure 8b(iii)**). Further, a pronounced difference was evident between normal skin and metastatic melanoma. Together, these results support a strong negative relationship between MTf and NDRG1 expression and confirm the importance of the MTf/NDRG1 regulatory axis in melanoma progression in patient samples.

To assess this further at the protein level, multiplex immunohistochemistry (mIHC) was performed using normal skin sections (*n* = 9), dysplastic nevi tissue microarrays (TMAs; *n* = 27), and treatment naïve primary (*n* = 31), and metastatic melanoma TMAs (*n* = 57; **Figure 8c**). In melanoma tumors, the pattern of NDRG1 (magneta fluorescence) distribution again clearly demonstrated the negative relationship with MTf (green fluorescence; **Figure 8c**). This is clearly shown in tumor regions where MTf is high and NDRG1 very low (indicated by a box with solid lines: **Figure 8c(i-iii)**), or in tumor regions where an interface exists between areas of high and low MTf expression that reciprocally demonstrate low and high NDRG1, respectively (denoted by a box with broken lines; **Figure 8c(i-iii)**). Areas with intermediate states exhibiting moderate expression of both MTf and NDRG1 were also observed in melanoma samples (**Figure 8c(i-iii), asterisk**). Examining the histoscores, most of the primary and metastatic melanoma samples were characterized by significantly increased MTf expression, compared to normal skin melanocytes (SOX10(+)) (**Figure 8d(i)**), whereas NDRG1 protein levels (**Figure 8d(ii)**) were markedly and significantly decreased.

The protein expression of MTf and NDRG1 (**Figure 8c(i,ii)**) strongly reflected the trends of gene expression at the mRNA level (**Figure 8b(i, ii)**). Indeed, the MTf histoscore gradually increased from the lowest values detected in normal melanocytes to metastatic melanoma samples exhibiting the highest expression of MTf protein (**Figure 8c(i)**). In contrast, the NDRG1 histoscore revealed a gradual decrease in levels with 7 out of 31 primary and 12 out of 57 metastatic tumors exhibiting completely negative NDRG1 expression (**Figure 8d(ii)**). Most impressively, the NDRG1/MTf ratio from the histoscores significantly correlated with melanoma progression (**Figure 8d(iii)**), which demonstrates the clinical significance of the MTf/NDRG1 axis in human patients.

## Discussion

MTf was one of the first melanoma tumor antigens that was well-characterized,^3^ yet its exact molecular function has remained mysterious.^8^ High MTf expression in melanoma cells and other tumor cells results in increased proliferation, migration, and tumorigenesis *in vivo*.^12, 14^ For the first time, the current study has demonstrated that MTf acts as a pro-oncogenic WNT agonist that down-regulates the metastasis suppressor and WNT antagonist, NDRG1, and *vice versa* in a Yin-Yang relationship (**Figure 8e**). The same intertwined association between these molecules was also evident using patient samples.

MTf is a known GPI-linked, plasma membrane protein.^63^ Surprisingly, cellular fractionation and confocal microscopy herein also demonstrated cytosolic and nuclear localization. Furthermore, *NDRG1* silencing increased nuclear, as well as cytoplasmic MTf, suggesting a pro-oncogenic role, as nuclear translocation is often related to oncogenic events.^27, 54–56, 64, 65^ Other membrane proteins can translocate from the plasma membrane to the nucleus by internalization *via* endocytosis, followed by fusion with the endoplasmic reticulum and transport to the nucleus *via* this system.^66–68^ MTf can be internalized into cells by mechanisms consistent with endocytosis and caveolae formation,^63, 69, 70^ but, there have been no previous reports of nuclear MTf expression. Other GPI-linked proteins, such as the folate receptor-α, can undergo nuclear translocation and act as a transcription factor.^71^ Considering this, another transferrin homologue, lactoferrin,^11^ demonstrates nuclear localization and potential transcriptional activity.^64, 65^ Hence, the nuclear localization of MTf may suggest a transcriptional role.^1, 4–7^

Considering the mechanism of action of MTf, it markedly down-regulated NDRG1 (**Figure 8e**), which inhibits the WNT pathway *via* its effect on β-catenin expression and distribution.^23, 27, 30^ In fact, NDRG1 promoted plasma membrane β-catenin localization and inhibited the EMT, while *NDRG1* silencing induced β-catenin nuclear translocation followed by activation of oncogenic targets *e.g.*, cyclin D1.^27^ Considering this, up-regulation of MTf, which decreases NDRG1 expression, also increased nuclear β-catenin. In contrast, *MTf* silencing distinctly increased localization of β-catenin to the plasma membrane. In addition, the nuclear p-β-catenin (Ser552), which activates WNT signaling,^49^ was also increased by MTf overexpression, with silencing having an opposite effect. Activation of WNT signaling by MTf was confirmed by the up-regulation of c-Myc and cyclin D1 (**Figure 5a**). Further, *NDRG1* silencing in melanoma cells resulted in nuclear MTf translocation. Hence, the pro-oncogenic effects of MTf are due to NDRG1 down-regulation, leading to increased WNT signaling.

Importantly, c-Myc is a known pro-oncogenic transcription factor^72–76^ that can also transcriptionally repress target genes.^77^ One of these targets is *NDRG1*, with c-Myc repressing its expression at its core promoter.^36, 58^ The studies showed herein that upon MTf overexpression, *c-Myc* mRNA and protein increased *via* the activation of the WNT pathway, with c-Myc being well known to repress NDRG1 expression (**Figure 8e**). Moreover, upon *c-Myc* silencing in MTf overexpression clones, both NDRG1 and MTf expression increased (**Figure 7f**). These results indicate the Yin-Yang relationship between NDRG1 and MTf was disrupted after *c-Myc* silencing, suggesting c-Myc plays a key linking role in the inverse regulation of MTf and NDRG1 (**Figure 8e**). It is also notable that *c-Myc* silencing increased MTf expression, suggesting a possible negative feedback mechanism.

The inverse relationship between MTf and NDRG1 is important not only for understanding melanoma biology and generating a prognostic marker, but also for therapeutic development. As such, the potent NDRG1-inducing thiosemicarbazones, Dp44mT and DpC,^61^ potently up-regulated NDRG1 and down-regulated MTf in melanoma cells, with Dp44mT inhibiting SK-MEL-28 melanoma xenograft growth *in vivo* In conclusion, MTf has a pro-oncogenic role mediated through WNT signaling and its ability to down-regulate the metastasis suppressor, NDRG1. A distinct Yin-Yang relationship existed between MTf and NDRG1 that could be disrupted after *c-Myc* silencing, suggesting a critical regulatory role of c-Myc. Finally, the pro-oncogenic role of MTf and its Yin-Yang relationship with NDRG1 means it can be effectively targeted *via* potent NDRG1-inducing agents to inhibit melanoma growth.

## Methods

### Cell Culture

The human melanoma cell-types, SK-MEL-28 and SK-MEL-2, were obtained from the American Type Culture Collection (ATCC, Manassas, VA). The patient derived melanoma cells WM793 and 1205 LU were a generous gift from Assoc. Prof. Elin Gray (Edith Cowan University, Western Australia). Cell lines were authenticated by the provider and this was based on viability, recovery, growth, morphology, cytogenetic analysis, antigen expression, DNA profile and iso-enzymology. Cells were routinely examined for mycoplasma contamination using standard methods. Both SK-MEL-28 and SK-MEL-2 were cultured in Eagle’s Minimum Essential Medium (MEM; Gibco, VIC, Australia) supplemented with 10% fetal calf serum (FCS; Sigma-Aldrich; St. Louise, MO), penicillin streptomycin glutamine (100 μg/mL; Invitrogen), sodium pyruvate (1 mM; Invitrogen) and non-essential amino acids (0.1 mM; Invitrogen). Overexpjression of MTf was accomplished by stably transfecting SK-MEL-28 cells with human *MTf* cDNA or with the empty pCMV-Script^®^ (Stratagene, CA, USA) vector alone^14^. The hyper-expressing clones are termed, hIF and hIE, and their vector control as VA, and maintained in the media above in G418 (1000 μg/ml; Alexis Biochemicals, Switzerland).^14^

In terms of stable *MTf* down-regulation model, SK-MEL-28 cells were stably transfected with the expression vector, p*Silencer* 3.1-H1 neo (Ambion, Texas).^12^ These clones were referred to as the B1 and B2 clones, and as a control, the cells were transfected with the same vector (SCR) containing a scrambled, non-specific siRNA insert (pS-scrambled; Ambion).^12^ The B1 clone demonstrated superior silencing of *MTf* and was used in preference in most experiments. The same procedure was used to down-regulate *MTf* expression in the SK-Mel-2 cells and are referred to as ‘A3’ with an appropriate SCR control.^14^ These stably transfected cells were selected and maintained in G418 (1000 μg/ml; Alexis Biochemicals, Lausen, Switzerland).^14^

The pancreatic cancer cell line PANC-1 was a generous gift from Prof. Andrew Biankin (Garvan Institute, NSW, Australia) and was grown in Dulbecco’s Modified Eagle Medium (DMEM; Gibco). For NDRG1 overexpression, the cells were stably transfected with pCMV-taq2-Flag-NDRG1 (GenHunter, Nashville, TN, USA), namely N1 and N2 or the empty pCMV-taq2-FLAG (Stratagene, Santa Clara, CA, USA) vector (VC) to serve as a negative control.^24^ All transfected cells were selected using G418, as described above.

### Protein Extraction, Western Blot, and Antibodies

Total protein was extracted using standard procedures in our laboratory.^23^ Western analysis was performed as described previously.^78^ The primary and secondary antibodies used are outlined in **Supplementary Table 1**. β-actin was used as a protein-loading control.

### Cytoplasmic and Nuclear Fractionation

Cytoplasmic and nuclear fractions were prepared from cultured cells using NE-PER cytoplasmic and nuclear extraction reagents (Thermo Fisher Scientific, Waltham, MA). Fractionation was performed according to the kit instructions. The lysates were then supplemented with a 1× solution of protease inhibitor (Roche Diagnostics) and 1× solution of PhosSTOP.

### Gene Silencing

Two specific siRNAs for *NDRG1* were used, namely, siNDRG1 (Cat.#: s20336; Life Technologies) and siNDRG1II (Cat.#: s20334; Life Technologies). These were compared to non-targeting negative control siRNA (siControl; Life Technologies). The siRNA was reverse transiently transfected into SK-MEL-28 and SK-MEL-2 cells using Lipofectamine RNAiMAX^®^ (Life Technologies) following the manufacturer’s instructions and incubated for 72 h/37°C, respectively. siRNAs specific for *MTf* (siMTf; Life Technologies), *c-Myc* (si-c-Myc, Cat.#: 6341S; Cell Signaling) were compared to non-targeting negative control siRNA (siControl; Life Technologies). The siRNAs were reverse transiently transfected into SK-MEL-28 cells using Lipofectamine RNAiMAX^®^ (Life Technologies) following the manufacturer’s instructions and incubated for 72 h/37°C, respectively. Western blotting was then performed, as described previously.^78^

### Immunofluorescence and Confocal Microscopy

Immunofluorescence was performed as described.^23^ Images were captured using a Zeiss LSM 510 Meta Spectral confocal microscope, a ZEISS LSM 800 plus Airyscan Spectral confocal microscope, and an Olympus FV3000RS NIR scanning confocal microscope using a 60x or 63x objective. Raw images were analyzed using AxioVision (Carl Zeiss, Australia), ImageJ plot profile tool (National Institutes of Health, Maryland), and/or the JACoP co-localization plugin.^79^

### MTT Cellular Proliferation Assay

Cellular proliferation was determined using the 3-(4,5-dimethylthiazol-2-yl)-2,5-diphenyl tetrazolium (MTT; Sigma) assay and validated by viable cell counts.^80^

### SK-MEL-28 Melanoma Xenografts and Treatment with Dp44mT

All animal studies were approved by the University of Sydney Animal Ethics Committee. SK-MEL-28 cells (10^7^) were suspended in Matrigel (BD Biosciences, San Jose, CA) and injected s.c. into the right flanks of female BALB/c nu/nu mice (8–10 weeks of age). After engraftment, tumor size was measured by Vernier calipers with tumor volumes (mm^3^) being calculated.^81^ When tumor volumes reached 120 mm^3^, *i.v.* treatment began. Agents were dissolved in 15% propylene glycol/0.9% saline and injected *i.v.* over 5 consecutive days/week for 2 weeks. Control mice were treated with vehicle alone.

### RNA Extraction and Semi-Quantitative RT-PCR Analysis

Cells were seeded onto tissue culture plates in MEM media (Gibco) and after 24 h, media was removed, and cells were rinsed with sterile PBS. RNA was extracted using TRIzol (Invitrogen) following manufacturers’ instructions (Applied Biosystems, CA, USA). Primers were designed and checked for specificity using the National Center for Biotechnology Information BLAST search tool (http://www.ncbi.nlm.nih.gov/tools/primer-blast/). Real time PCR quantification of *MTf*, *NDRG1* and *c-Myc*, genes was carried out using standard procedures as described in^59^ with gene-specific primers **(Supplementary Table 2)**. *β-actin* was used as a loading control.

The following thermocycling optimization protocol was used: 56°C for 30 min, 94°C for 2 min and 15 s, 58°C/30s followed by the appropriate number of cycles for each primer at 94°C/15 s/cycle (**Supplementary Table 3**). For the elongation step, samples were heated at 68°C/5 min and finally cooled to 4°C. The following thermocycling optimization protocol was used: 56°C/30 min, 94°C/2 min and 15 s, 58°C/30 s. followed by the appropriate number of cycles for each primer at 94°C/15 sec per cycle (**Supplementary Table 3)**. For the elongation step, the samples were heated at 68°C/5 min and finally cooled to 4°C. Assessment of DNA fragment length was performed by electrophoretic separation in a 0.4% agarose gel with DNA ladder (Promega, WI, USA).

### Tissue Microarrays and Tissue Sections

Tissue microarrays (TMAs) constructed from formalin-fixed paraffin-embedded (FFPE) lesions, ranging from compound naevus to distant melanoma metastases, and normal skin tissues were retrieved from the Department of Tissue Pathology and Diagnostic Oncology at the Royal Prince Alfred Hospital, Australia. Studies using human tissues were approved by the Human Research Ethics Committees of the University of Newcastle and Royal Prince Alfred Hospital, Australia.

### Quantitative Multiplex Immunohistochemistry (mIHC)

All immunofluorescence staining was carried out on TMAs and normal skin tissue sections using an Autostainer Plus (Dako, Agilent Technologies) with appropriate positive and negative controls. Opal Multiplex IHC Assay kit (PerkinElmer, USA) was used as per the manufacturer’s protocol. Briefly, FFPE tissue sections were first deparaffinised and rehydrated using xylene and ethanol. Heat-induced antigen retrieval (AR) was performed in a Decloaking Chamber (Biocare) in pH 6 AR buffer at 95°C/20 min. Endogenous peroxidase activity was quenched with 3% hydrogen peroxide for 5 min. Individual primary antibodies targeting MTf (1:100; HPA004880, Sigma), NDRG1 (1:800; D8G9 XP, Cell Signaling Technology), and SOX10 (1:1500; ab212843, Abcam) were incubated for 30 min. Primary antibodies were detected using Opal Polymer HRP (Perkin Elmer) and visualized using Tyramide Signal Amplification for 10 min (Opal 7-Colour IHC, Perkin Elmer). Sections were stained with DAPI for 5 min and mounted using ProLong® Diamond. Imaging was performed using the Vectra 3 multispectral slide scanner in conjunction with Vectra 3.3 and Phenochart 1.0.4 software (PerkinElmer). Images were unmixed in inForm 2.2.0 and a selection of 15 representative original multi-spectral images was used to train the single-cell separation algorithm in HALO™ Image Analysis software (Indica Labs, melanocytic cells identified based on SOX10 expression). All the settings applied to the training images were saved within an algorithm to allow batch analysis of multiple original multi-spectral images of the individual TMA cores.

### Statistical Analysis

Data were compared using Student’s *t*-test. Gene Expression Omnibus (GEO) and mIHC data were compared using one-way ANOVA and the Mann-Whitney U test, respectively. Data were considered statistically significant when *p* < 0.05. Results are expressed as mean ± SD or mean ± SEM.

## Acknowledgments

D.R.R. appreciates NHMRC Senior Principal Research Fellowships [APP1159596; APP1062607], NHMRC Project Grant [APP1144456] and NHMRC/PdCCRS Grant [APP1146599]. Z.K. is grateful for a University of Sydney (USYD) Bridging Fellowship, a National Health and Medical Research Council (NHMRC) RD Wright Fellowship [1140447], a Cancer Institute New South Wales (CINSW) Career Development Fellowship [CDF171126] and USYD Equity Fellowship.

## Contributions

D.R.R, J.P., Z.K and D.J.L. conceived and designed the study. J.P. , M.G.A. and D.R.R. oversaw the execution of data analysis and produced the output figures and tables. M.L.H and J.S. participated in study optimization and data analysis. J.S. oversaw the study design, execution and data analysis of Quantitative Multiplex Immunohistochemistry. K.C.P., R.A., S.C., D.H. and G.G.A. contributed to western blotting and figure formatting. R.S. and J.W. retrieved human tissue samples for tissue microarray construction and validated the testing protocol. J.P. and M.J.S created the NDRG1 over-expressing SK-MEL-28 cells and oversaw the study design, execution, and data analysis. K.L. and M.W. performed the MTT and *in vivo* study, respectively and J.P. performed the data analysis. D.R.R., J.P. and D.J.L. wrote the initial manuscript, and all authors contributed to subsequent revisions. D.R.R. and J.P. had final responsibility for the decision to submit for publication after consultation with all authors.

## Figure Legends

## Supplemental Figure Legends

**Supplemental Figure 1. Silencing MTf up-regulates NDRG1, with the Yin-Yang relationship between MTf and NDRG1 being also present in melanoma patient-derived cells.** (a) Silencing *MTf* expression using a stable siMTf clone (B2) of SK-MEL-28 cells results in the up-regulation of NDRG1 *versus* the scrambled (SCR) control, as demonstrated by western blotting. (b) Higher levels of MTf correlate with decreased NDRG1 expression in patient derived melanoma cells as demonstrated by western blotting. Both 1205 LU and 451 LU are the metastatic forms of the primary tumor cells, WM793 and WM164, respectively. Results in (a, b) are representative blots and the densitometry is presented as mean ± SD (3 experiments). \**p*<0.05 \*\**p*<0.01 \*\*\**p*<0.001 relative to the SCR or VA control, respectively.

**Supplemental Figure 2. MTf expression decreases p-NDRG1 levels at Ser330 and Thr346 in SK-MEL-28 cells. (a)** Relative to VA SK-MEL-28 cells, the MTf overexpressing clones hIF and hIE down-regulate p-NDRG1 at Ser330 and Thr346, as demonstrated by western blotting. **(b)** Cells transiently transfected with either *MTf* siRNA or the negative control siRNA (NC). **(c)** Cytoplasmic (C) and nuclear (N) fractions of VA, hIF and hIE cells demonstrate decreased p-NDRG1 at Ser330 and Thr346 levels, with fraction purity being monitored by probing for histone deacetylase (HDAC; nucleus) and glyceraldehyde-3-phosphate dehydrogenase (GAPDH; cytosol). Results in **(a-c)** are representative blots and the densitometry is presented as mean ± SD (3 experiments). \**p*<0.05 \*\**p*<0.01 \*\*\**p*<0.001 relative to the respective control. (**d,e**) Confocal immunofluorescence microscopy of the VA, hIF and hIE MTf overexpression model as well as NC and siMTf cell models examining: **(d)** p-NDRG1 Ser330 and **(e)** p-NDRG1 Thr346 intensity and their subcellular localization. The software ImageJ and its analytical tool, co-masking analysis, were used to examine p-NDRG1 Ser330 and Thr346 levels (total, cytoplasmic, and nuclear). Results are mean ± SD (3 experiments). \**p*<0.05, \*\**p*<0.01, \*\*\**p*<0.001 relative to the respective control. ImageJ analysis utilized a total of 12-22 cells over 3 experiments. All images were taken using an 63x objective and the scale bar in the bottom left-hand corner of the first image in **(d)** and **(e)** represents 30 μm and is the same across all images.

**Supplemental Figure 3. Silencing NDRG1 with siNDRG1 II in SK-MEL-28 melanoma cells increases MTf expression, while NDRG1 overexpression down-regulates MTf.** (a) Transient silencing of *NDRG1* with an additional *NDRG1* siRNA (siNDRG1 II) in SK-MEL-28 cells results in up-regulation of MTf, β-catenin and p-AKT *versus* the negative control (NC) siRNA, as shown by western blotting. (bi, ii) Immunofluorescence microscopy demonstrating up-regulation of NDRG1 and down-regulation of MTf in the PANC-1 stable overexpressing NDRG1 clone (N1) relative to vector control (VC) cells. Images were taken using a 42× objective with the scale bar in the first image in (bi & ii) representing 30 μm and is the same across all images. Quantitative analysis of NDRG1 and MTf intensity was performed using ImageJ with results being mean ± SD (3 experiments).

**Supplemental Figure 4. (a) Transfection of SK-MEL-28 melanoma cells with increasing levels of an NDRG1 expression plasmid increases NDRG1 and decreases MTf and LRP6 expression.** SK-MEL-28 cells were transiently transfected with increasing levels of the empty expression vector or this vector containing NDRG1 cDNA (0.375, 0.75, or 1.25 µg). Western analysis was then performed to examine LRP6, MTf, and NDRG1 expression relative to the β-actin loading control. (b) MTf hyperexpression increases the half-life of LRP6 protein levels in SK-MEL-28 cells. The VA control and MTf hyperexpression clone (hIF) were preincubated with cycloheximide (15 mg/ml) for 1 h/37°C to inhibit protein synthesis. After this 1 h preincubation, cells were harvested as the first timepoint (0 h), or the incubation with cycloheximide continued and cells harvested after 2-, 4-, 8-, 10-, and 12-h/37°C. Western analysis examined LRP6, MTf, and NDRG1 expression with β-actin being used as a protein loading control. Results are mean ± SD (3 experiments).

**Supplemental Figure 5.** C**o-immunoprecipitation and confocal microscopy demonstrates that LRP6 and β-catenin associate with MTf**. **(a,b)** LRP6 was co-immunoprecipitated and western blotting performed to examine LRP6 and MTf in: **(a)** MTf overexpressing hIF and hIE clones, *versus* the VA control, and **(b)** the stable *MTf* silenced B1 clones *versus* the SCR control, **(c, d)** β-catenin was co-immunoprecipitated and western blotting performed examining β-catenin and MTf in: **(c)** MTf overexpressing hIF and hIE clones, *versus* the VA control; and **(d)** the stable *MTf* silenced B1 clones *versus* the SCR control. **(e, f)** Confocal microscopy demonstrating co-localization between **(e)** LRP6 and MTf and **(f)** β-catenin and MTf in MTf overexpressing hIF and hIE clones, *versus* the VA control; and the stable *MTf* silenced B1 clones *versus* the SCR control. The ImageJ co-localization analysis tool was implemented and utilized 11-52 cells over 3 experiments. Scale bar = 3.8 μm. Results are mean ± SD (3 experiments). \**p*<0.05, \*\**p*<0.01, \*\*\**p*<0.001 *versus* the control.

**Supplemental Figure 6. MTf overexpression in SK-MEL-28 melanoma cells increases total and nuclear p-LRP6 (Ser1490), while silencing *MTf* has the opposite effect. (a)** Confocal microscopy was performed in VA relative to hIE and hIF MTf overexpressing cells and *MTf* silenced B1 cells *versus* SCR with p-LRP6 (Ser1490) localization examined using ImageJ plot profile analysis (bi-v) and co-masking analysis (c). Results are mean ± SD (3 experiments). \**p*<0.05, \*\**p*<0.01, \*\*\**p*<0.001 relative to the respective control. ImageJ analysis utilized a total of 15-41 cells over 3 experiments. All images were taken using an 63x objective and the scale bar in the bottom left-hand corner of the first image in (a) represents 30 μm and is the same across all images. The white line that crosses the cell body in the merged image displays intensities of different channels in the plot profile analysis.

**Supplemental Figure 7. Confocal immunofluorescence microscopy demonstrates that MTf overexpression in SK-MEL-28 cells increases total, nuclear, and cytoplasmic cyclin D1 levels, while *MTf* silencing results in the opposite response. Silencing *MTf* expression up-regulates total β-catenin and down-regulates c-myc and cyclin D1 levels. (a)** Confocal immunofluorescence microscopy and co-Masking Analysis *via* ImageJ demonstrates that cyclin D1 expression is increased the total, cytoplasmic and nuclear compartments of MTf overexpressing vector-transfected clones (hIF and hIE) relative to the vector transfected control cells (VA), while transient *MTf* silencing in SK-MEL-28 cells decreases the total, cytoplasmic and nuclear cyclin D1 levels *versus* the negative control (NC). (b) Silencing *MTf* expression using a stable siMTf clone (B1) of SK-MEL-28 cells results in the up-regulation of total β-catenin, and down-regulation of c-Myc and cyclin D1 *versus* the scrambled (SCR) control, as demonstrated by western blotting. (c) Silencing *MTf* expression using a stable siMTf clone (B2) of SK-MEL-28 cells results in down-regulation of c-Myc and cyclin D1 *versus* the scrambled (SCR) control, as demonstrated by western blotting. Results are mean ± SD (3 experiments). \*\**p*<0.01, \*\*\**p*<0.001 *versus* the relevant control. ImageJ analysis utilized a total of 11-21 cells over 3 experiments. All images were taken using an 63x objective and the scale bar in the first image in (a) represents 30 μm and is the same across all images.

**Supplemental Figure 8.** M**T**f **overexpression in SK-MEL-28 melanoma cells increases p-β-catenin (Ser552) in the nucleus and cytoplasm**. **(a-c)** Confocal immunofluorescence microscopy examining the subcellular localization of p-β-catenin S552 in SK-MEL-28 MTf overexpressing hIE and hIF clones *versus* VA control cells, or after siMTf treatment *versus* the NC. ImageJ plot profile analysis (**b**) and co-masking analysis (**c**) of the images in (**a**). The white line that crosses the cell body (**a**) results in the intensities of the different channels displayed in the plot profile. ImageJ analysis utilized 27-55 cells over 3 experiments. Scale bar = 30 μm. Results are mean ± SD (3 experiments). \**p*<0.05, \*\**p*<0.01, \*\*\**p*<0.001 relative to the respective control.

**Supplemental Figure 9.** M**T**f **overexpression enhances SK-MEL-28 proliferation, while stable *MTf* silencing has the opposite effect. (a)** Relative to VA control cells, the MTf overexpressing hIF and hIE SK-MEL-28 melanoma cells increased proliferation, as judged by MTT assays and validated by viable cell counts using Trypan blue. **(b)** Stable silencing of *MTf* in SK-MEL-28 cells (clones B1 and B2) decreases proliferation, as judged by MTT assays and validated by viable cell counts. Results are mean ± SD (3 experiments). \**p*<0.05, \*\**p*<0.01, \*\*\**p*<0.001 relative to the respective control.

**Supplemental Figure 10. DpC and Dp44mT up-regulates NDRG1, while down-regulating MTf and cyclin D1 expression SK-MEL-28 melanoma cells and Dp44mT effectively inhibits SK-MEL-28 melanoma tumor growth *in vivo*.** Western blot analysis of: (a) MTf, NDRG1 and cyclin D1 expression after treatment with DpC (1 μM, 5 μM, 10 μM) after an incubation of 24 h/37°C and compared to their control in VA. Results in (b,d) are representative blots and the densitometry is presented as mean ± SD (3 experiments). \**p*<0.05 \*\**p*<0.01 \*\*\**p*<0.001 relative to the control lysate.

**Supplemental Figure 11. Confocal immunofluorescence microscopy demonstrates that in SK-MEL-28 cells Dp44mT up-regulates NDRG1 increasing it co-localization with LRP6 (a-d), but down-regulates MTf and decreases LRP6 expression and its co-localization with LRP6 (e-h).** Confocal immunofluorescence microscopy and the ImageJ analysis tool and JACoP colocalization plugin were used to investigate intensity, co-localization between NDRG1 and LRP6 after a 24 h incubation with control medium, or this medium containing Dp44mT (5 µM) Results are mean ± SD (3 experiments). \*\**p*<0.01; \*\*\**p*<0.001 *versus* the relevant control. ImageJ analysis utilized a total of 18-21 cells over 3 experiments. All images were taken using an 60x objective and the scale bar in the first image in (a) represents 30 μm and is the same across all images except the 3x magnified images where it is 10 µm.

**Supplemental Figure 12.** D**p**44mT **effectively inhibits SK-MEL-28 melanoma tumor growth *in vivo*.** Administration of Dp44mT (*i.v.*, 0.625, 0.65 mg/kg, 0.675, and 0.7 mg/kg *per* day/5 days/week) of Balb c nu/nu mice significantly inhibits net tumor growth of SK-MEL-28 melanoma tumor xenografts over a treatment period of 2 weeks. Each dose represents mean ± SEM from 4-6 mice. Results are presented as mean ± SD (3 experiments). \**p*<0.05; \*\**p*<0.01; \*\*\**p*<0.001 relative to the vehicle control.

**Supplementary Table 1:** Antibodies used.

| **Antibody** | **Company** | **Catalog#** | **Dilution** |
| --- | --- | --- | --- |
| MTf | Sigma Aldrich, St. Louis, MO | HPA004880 | 1:1000, IF: 1:300 |
| NDRG1 | Abcam, Cambridge, MA, USA | ab37897 | 1:2000 |
| NDRG1 XP | Cell Signaling Technology,<br>Danvers, MA, USA | 9485S | IF: 1:150 |
| p-NDRG1 S330 | Cell Signaling Technology,<br>Danvers, MA, USA | 11899S | 1:2000, IF: 1:200 |
| p-NDRG1 S346 | Cell Signaling Technology,<br>Danvers, MA, USA | 5482S | 1:3000, IF: 1:250 |
| LRP6 | Cell Signaling Technology,<br>Danvers, MA, USA | 2560S | 1:1000, IF: 1:250 |
| LRP6 | Abcam, Cambridge, MA, USA | ab75358 | 1:1000, IF: 1:250 |
| p-LRP6 S1490 | Cell Signaling Technology,<br>Danvers, MA, USA | 2568S | 1:500, IF: 1:100 |
| β-catenin | Abcam, Cambridge, MA, USA | ab32572 | 1:10000 |
| β-catenin | Abcam, Cambridge, MA, USA | ab22656 | 1:250 |
| c-Myc | Cell Signaling Technology,<br>Danvers, MA, USA | 5605S | 1:1000, IF: 1:300 |
| cyclin D1 | Cell Signaling Technology,<br>Danvers, MA, USA | 2922S | 1:1000, IF: 1:300 |
| HDAC | Cell Signaling Technology,<br>Danvers, MA, USA | 2062S | 1:1000 |
| GAPDH | Cell Signaling Technology,<br>Danvers, MA, USA | 5174S | 1:10000 |
| β-actin | Sigma Aldrich, St. Louis, MO | A2228-100UL | 1:10000 |
| anti-goat secondary<br>antibody | Sigma-Aldrich, St. Louis, MO | A5420 | 1:10000 |
| anti-rabbit<br>secondary antibody | Sigma-Aldrich, St. Louis, MO | AP307P | 1:10000 |
| anti-mouse<br>secondary antibody | Sigma-Aldrich, St. Louis, MO | M8645 | 1:10000 |
| Goat Anti-Mouse<br>IgG Polyclonal<br>Antibody (IRDye<br>800CW) | Li-COR Biotechnology, Nebraska,<br>USA | 926-3221 | 1:10000 |
| Goat Anti-Rabbit<br>IgG Polyclonal<br>Antibody (IRDye<br>680LT) | Li-COR Biotechnology, Nebraska,<br>USA | 926-68021 | 1:10000 |
| Donkey Anti-Goat<br>IgG Polyclonal<br>Antibody (IRDye<br>680LT) | Li-COR Biotechnology, Nebraska,<br>USA | 926-68024 | 1:10000 |

**Supplementary Table 2:**
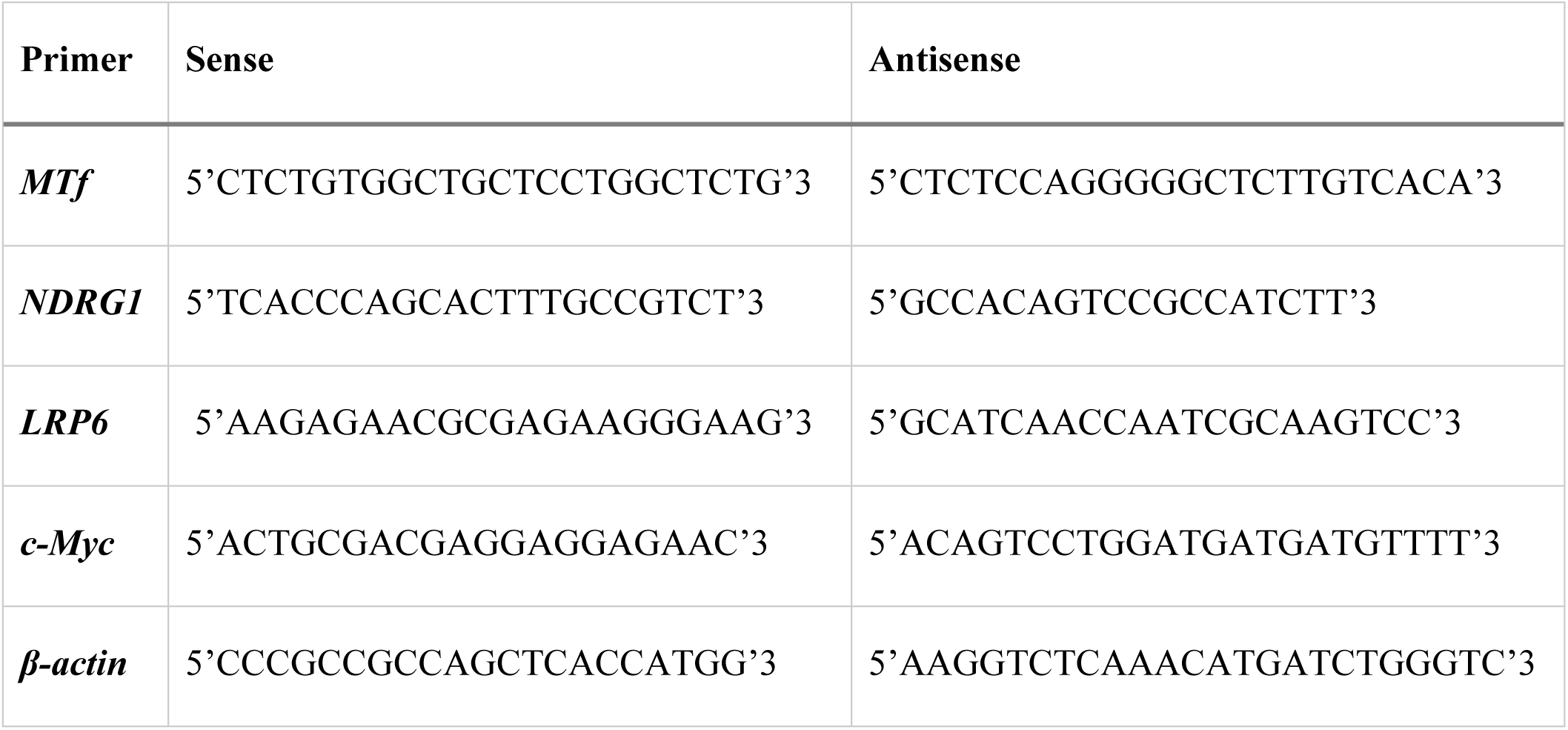
Sequences for the sense and antisense primers used.

**Supplementary Table 3:** The PCR cycle numbers used to detect *MTf*, *NDRG1*, *c-Myc*, and β-actin.

| **Primer** | *MTf* | *NDRG1* | *LRP6* | *c-Myc* | *β-actin* |
| --- | --- | --- | --- | --- | --- |
| **Cycles** | 25 | 27 | 29 | 27 | 24 |

