## Supplemental Figures for "Melanotransferrin Functions as a Pro-Oncogenic WNT Agonist: A Yin-Yang Relationship in Melanoma with the WNT Antagonist and Metastasis Suppressor, NDRG1"

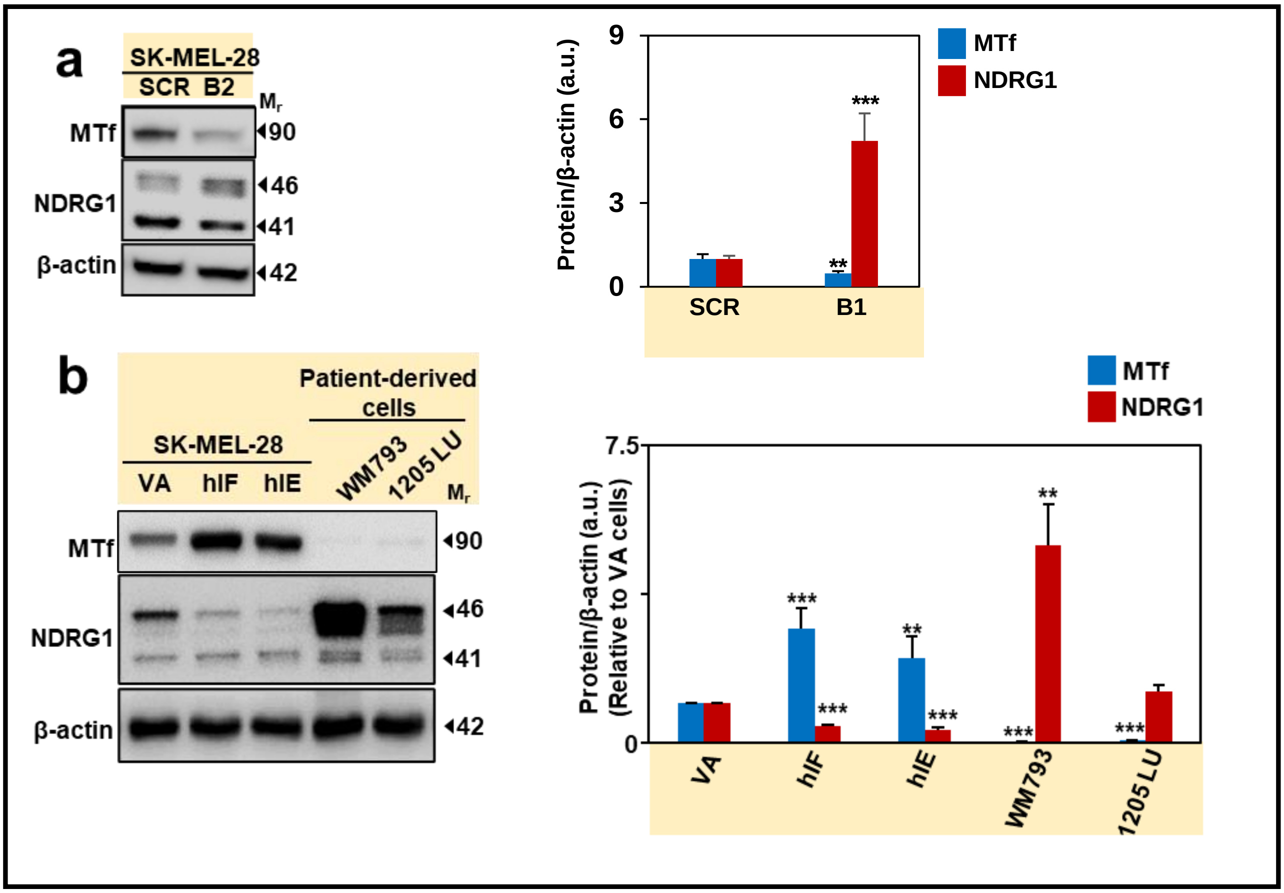
a
SK-MEL-28
SCR B2
Mr
MTf
NDRG1
β-actin
90
46
41
42
Protein/β-actin (a.u.)
9
6
3
0
SCR
B1
MTf
NDRG1
**
***
b
SK-MEL-28
VA hIF hIE
Patient-derived cells
WM793
1205 LU
Mr
MTf
NDRG1
β-actin
90
46
41
42
Protein/β-actin (a.u.)
(Relative to VA cells)
7.5
0
VA
hIF
hIE
WM793
1205 LU
MTf
NDRG1

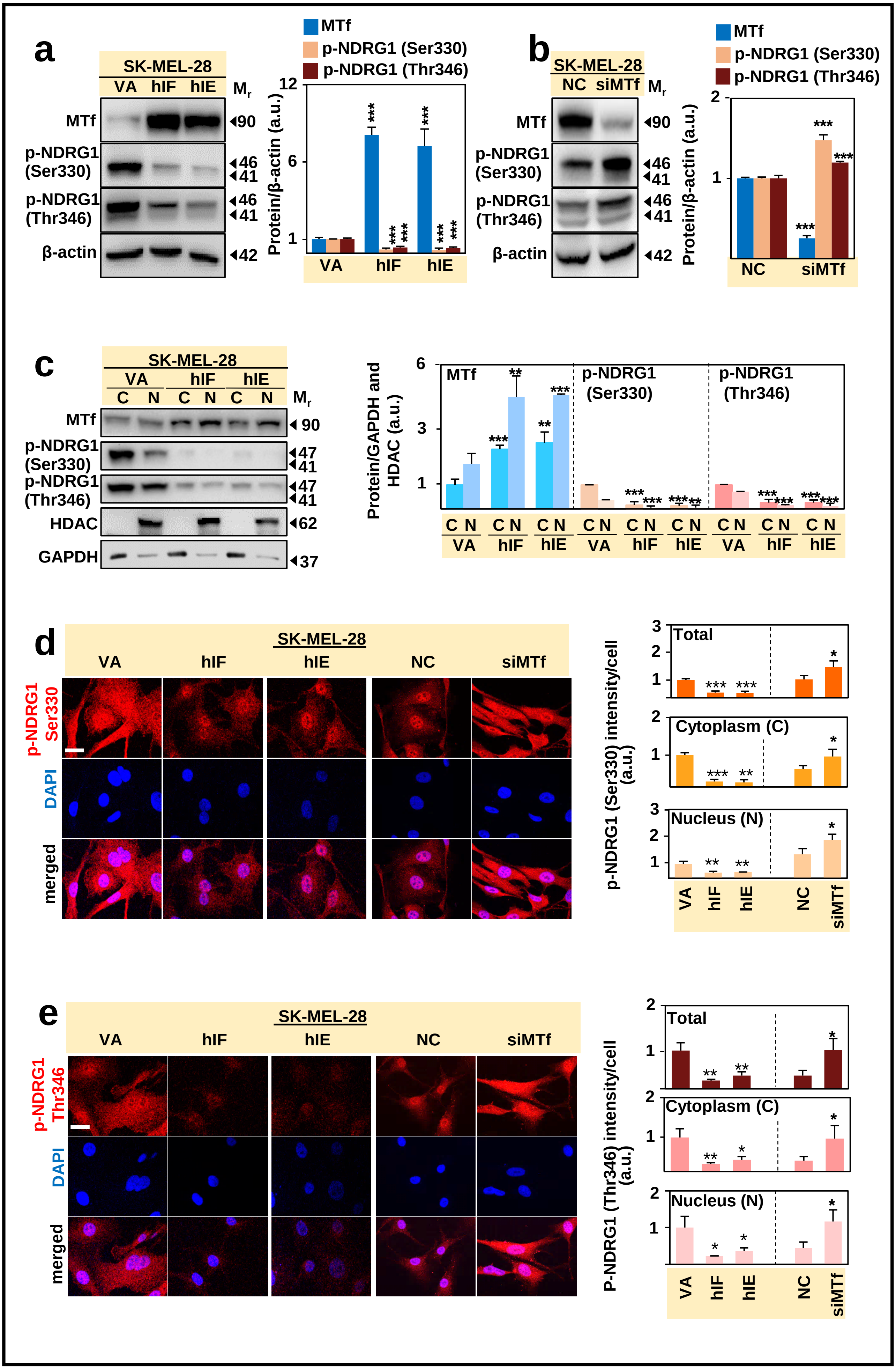


Supplemental Figure 2

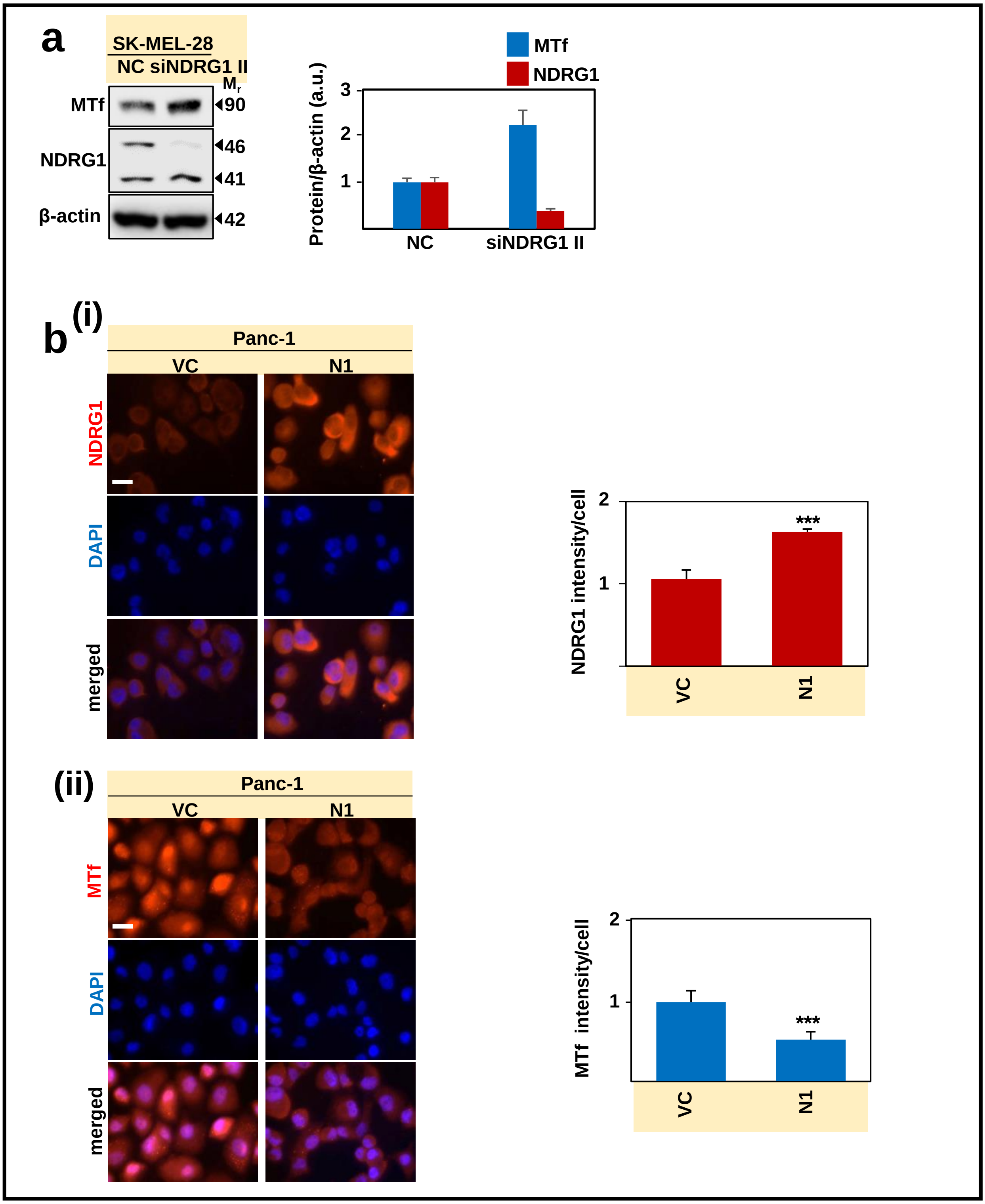


Supplemental Figure 3

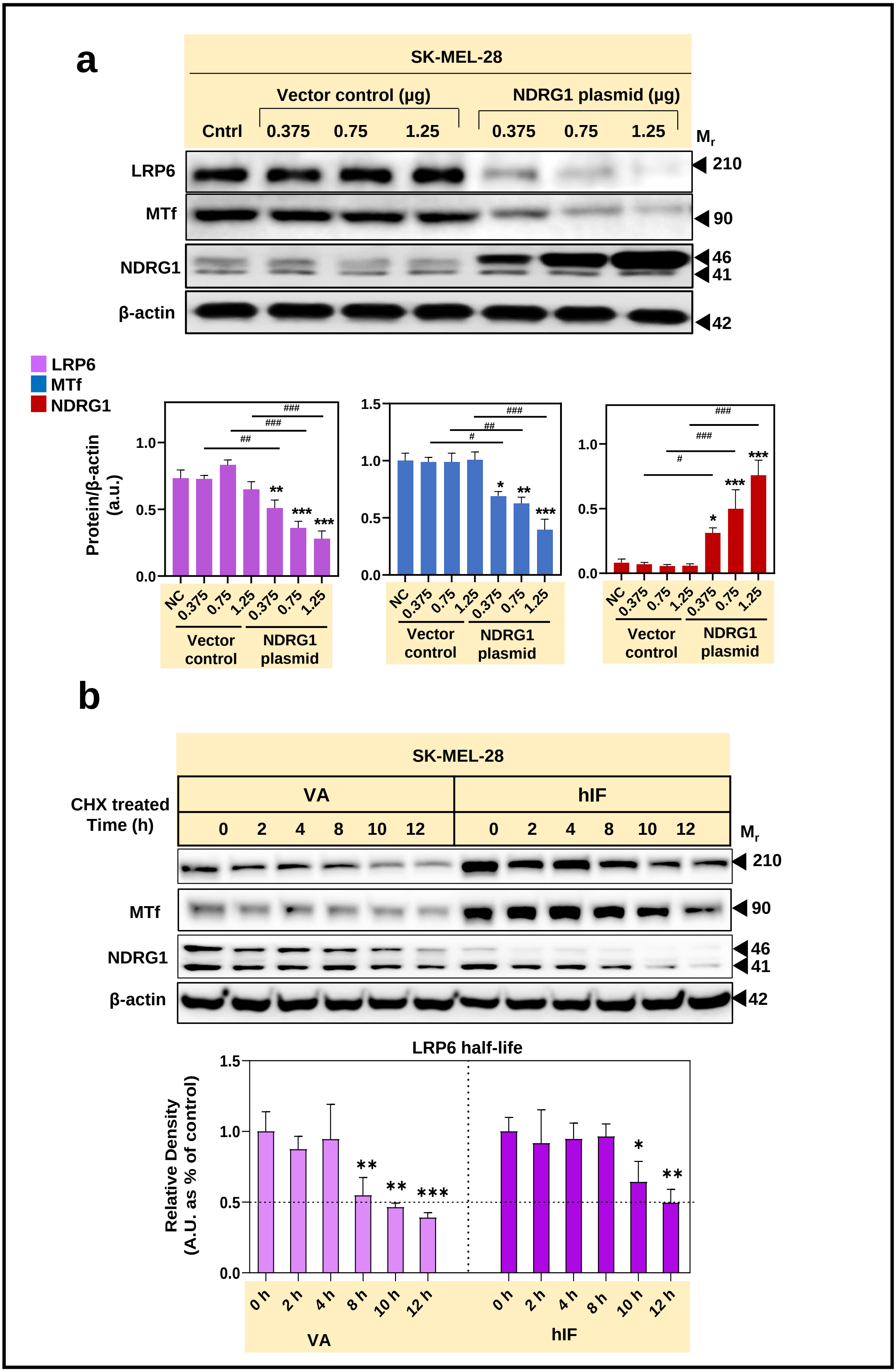


Supplemental Figure 4

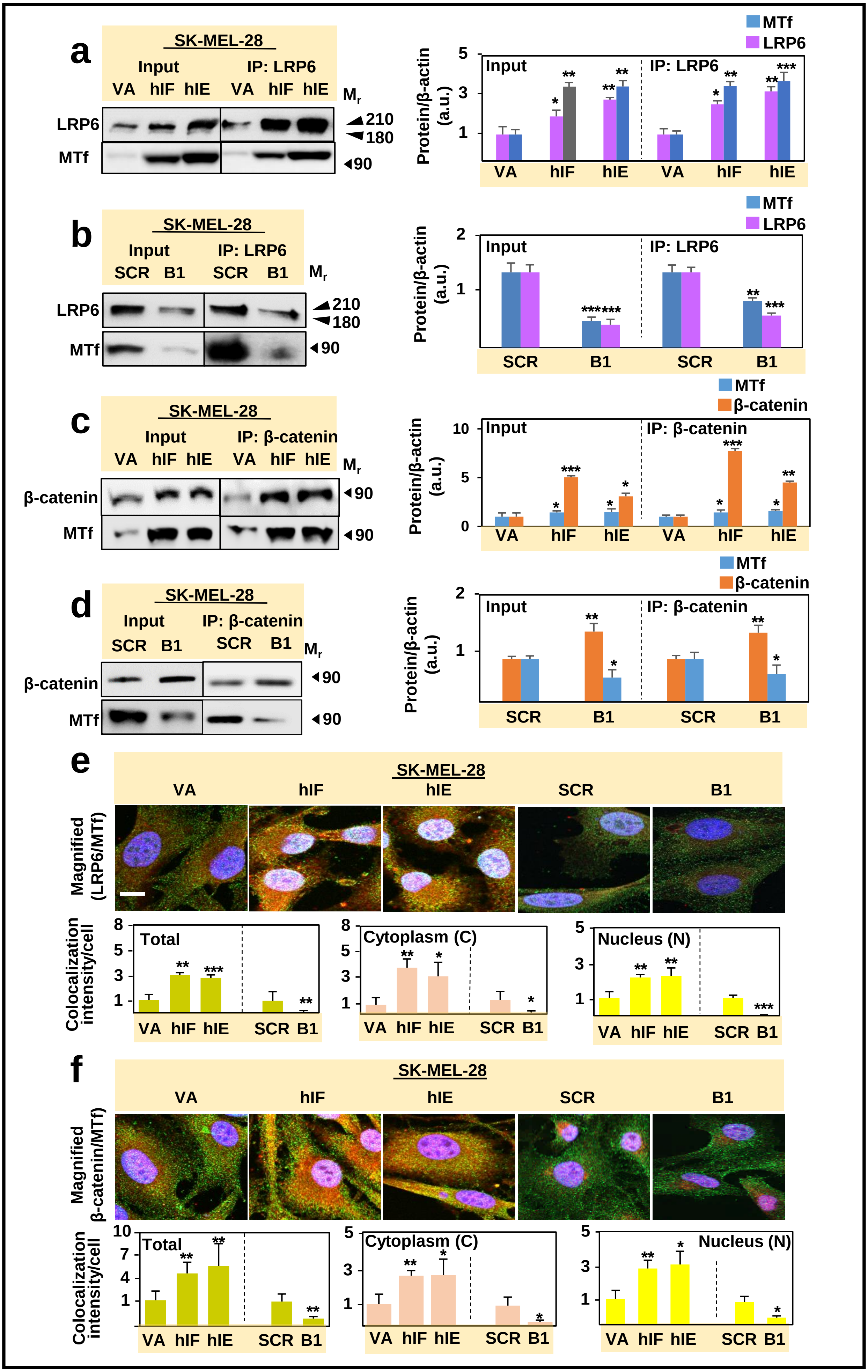


Supplemental Figure 5

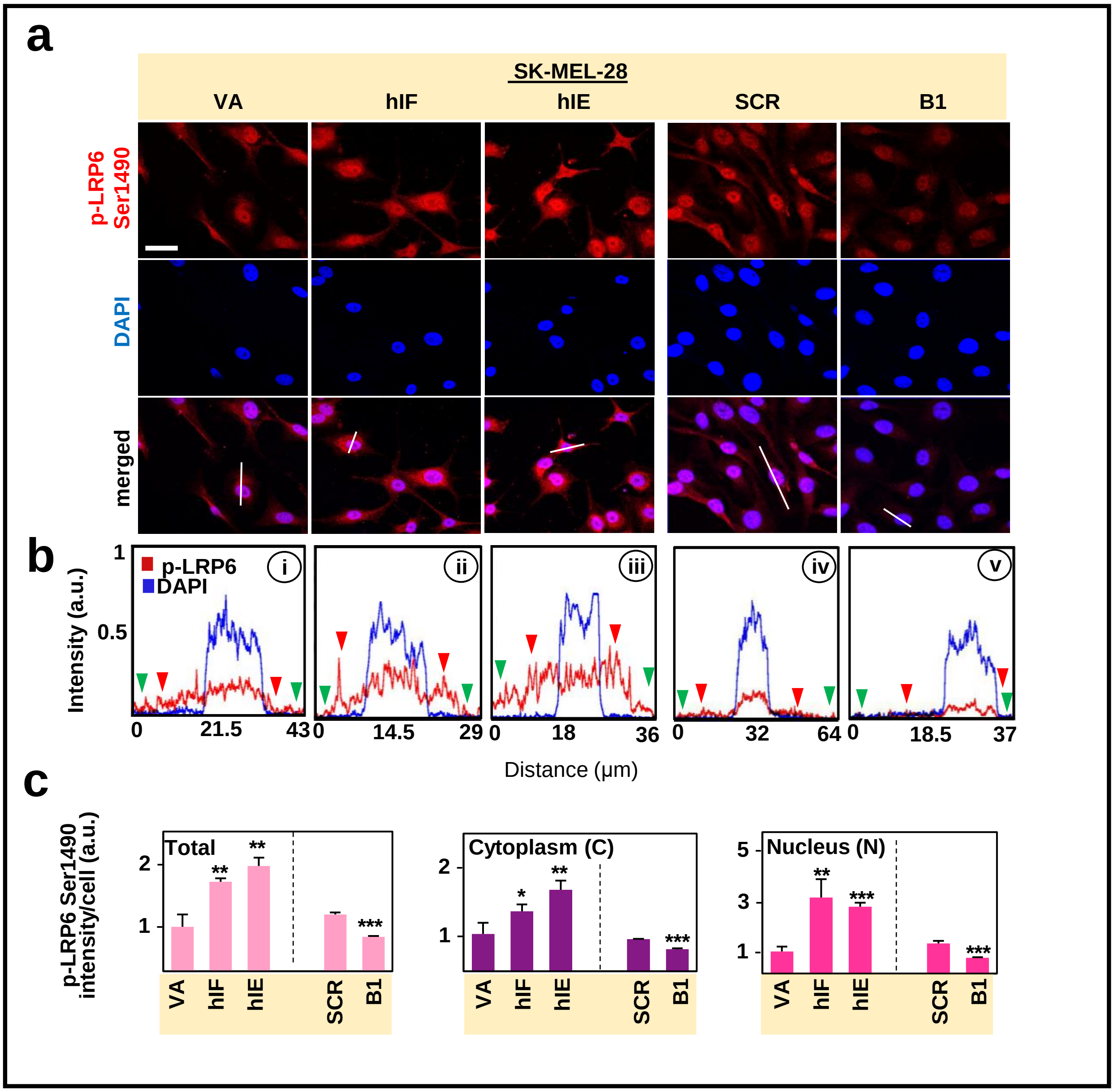


**Supplemental Figure 6**

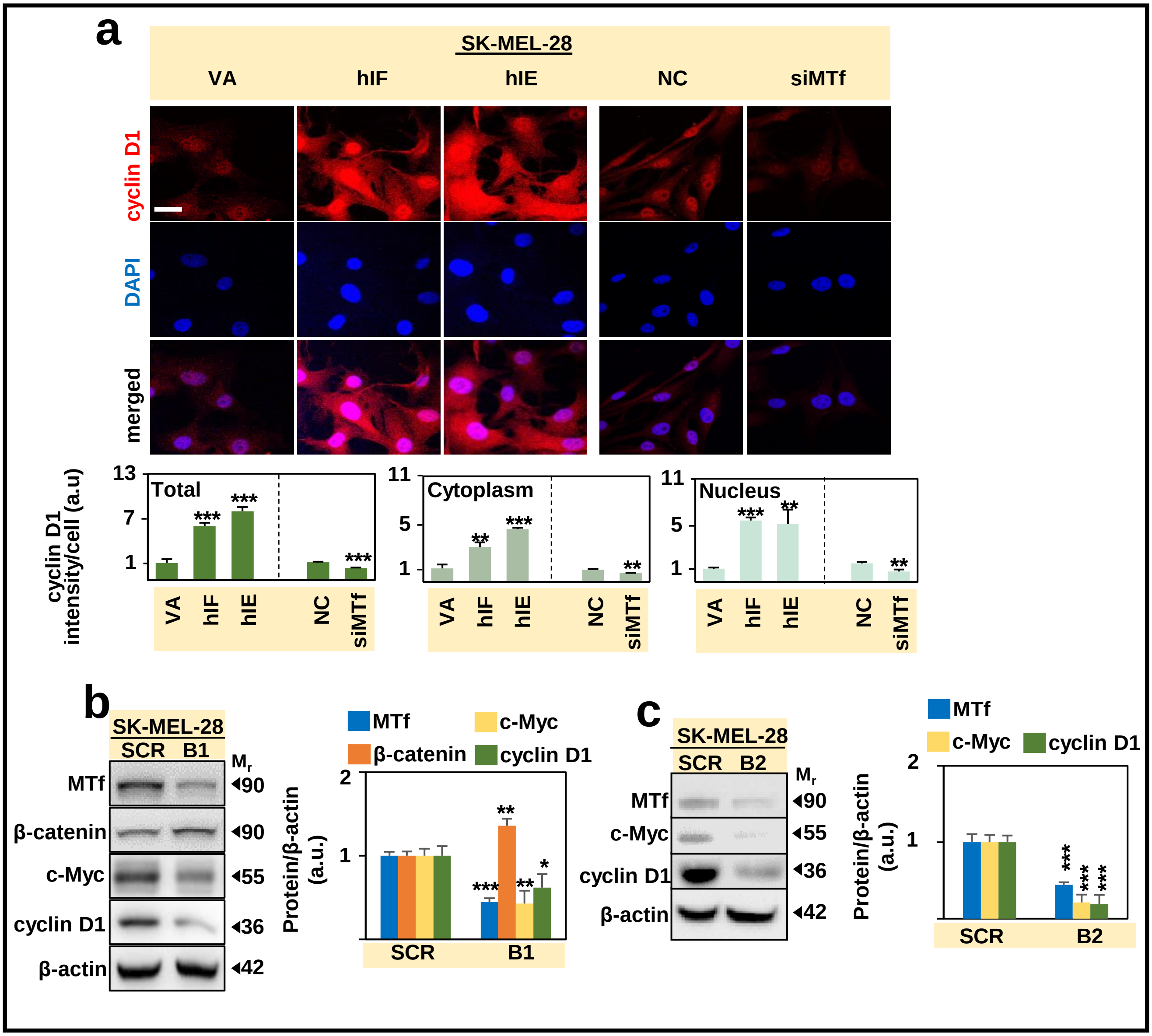


Supplemental Figure 7

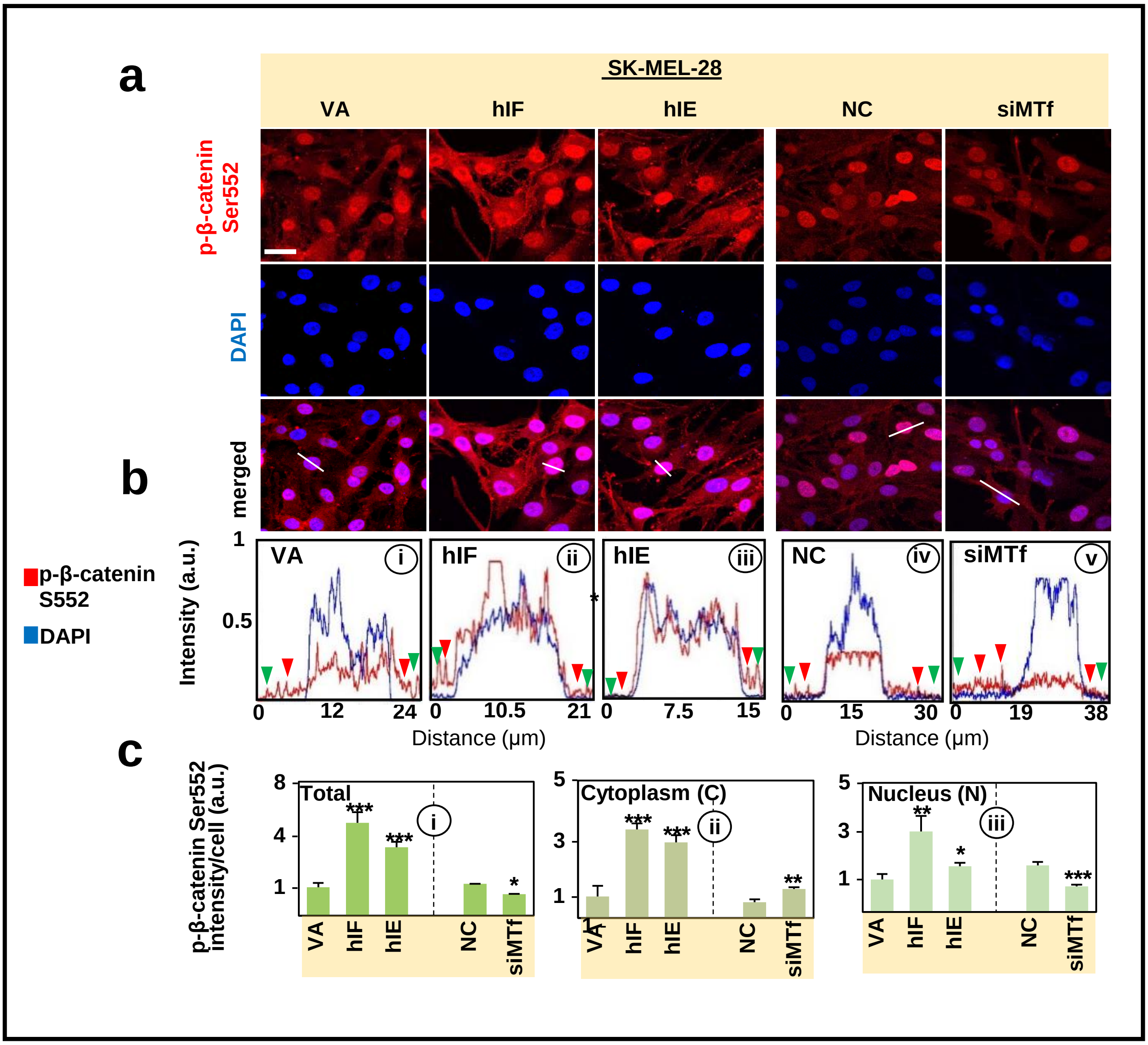


Supplemental Figure 8

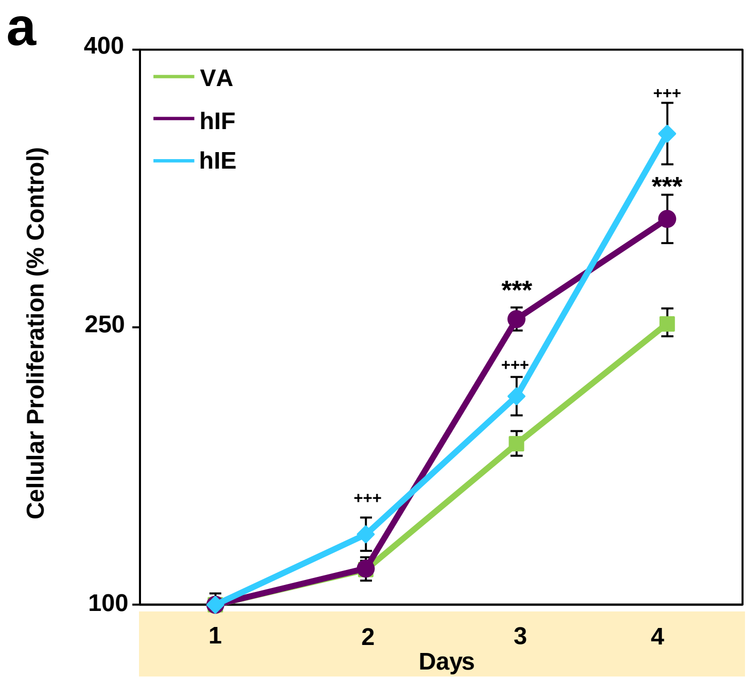


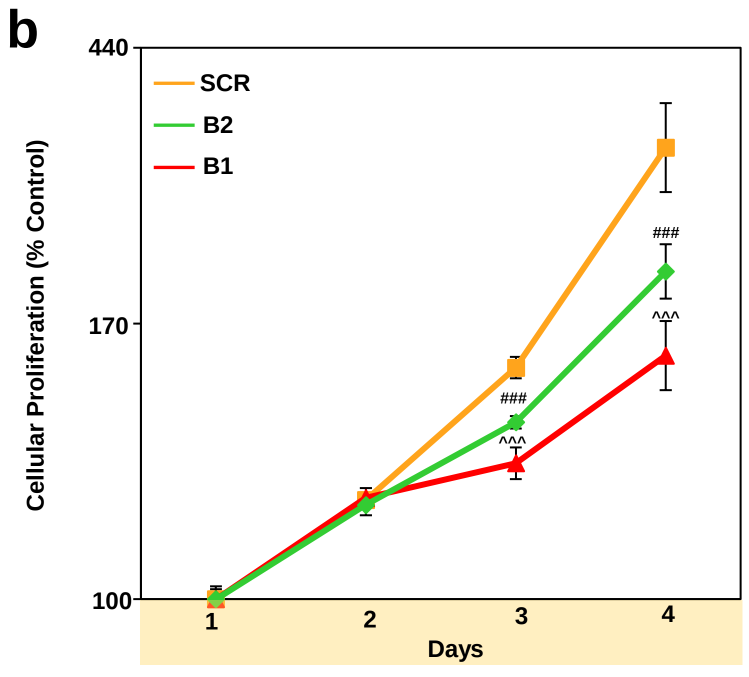


Supplemental Figure 9

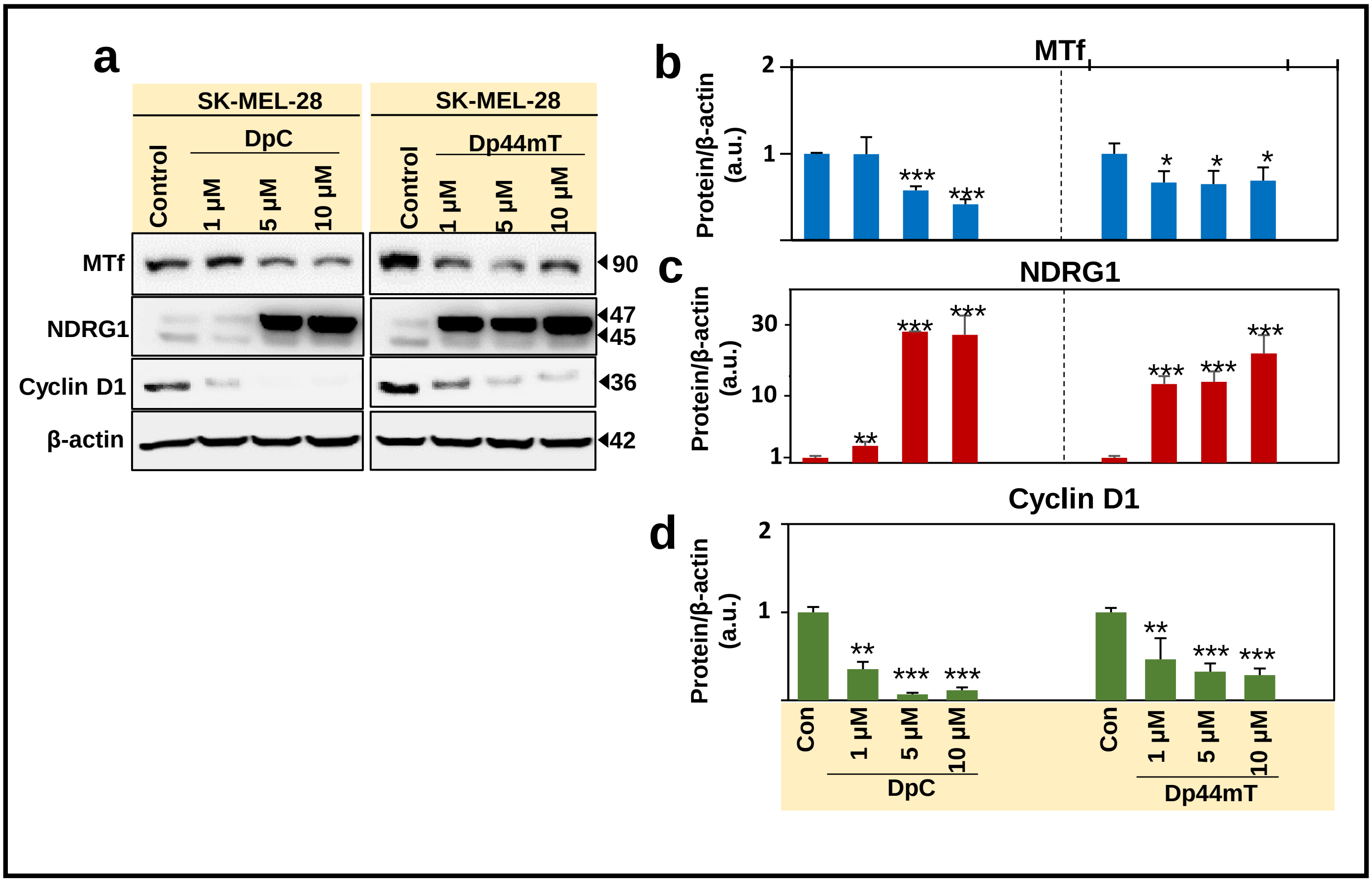


Supplemental Figure 10

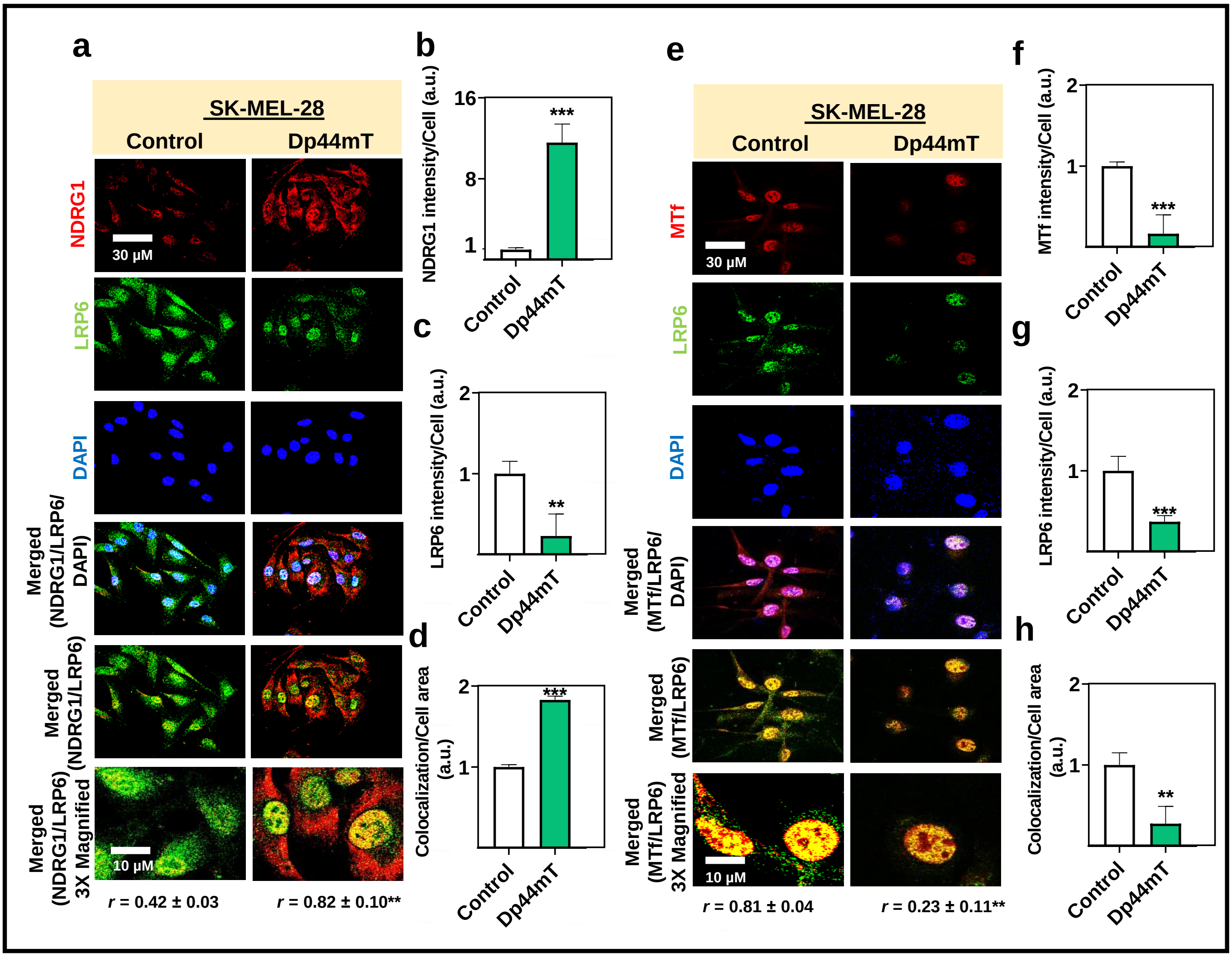


**Supplemental Figure 11**

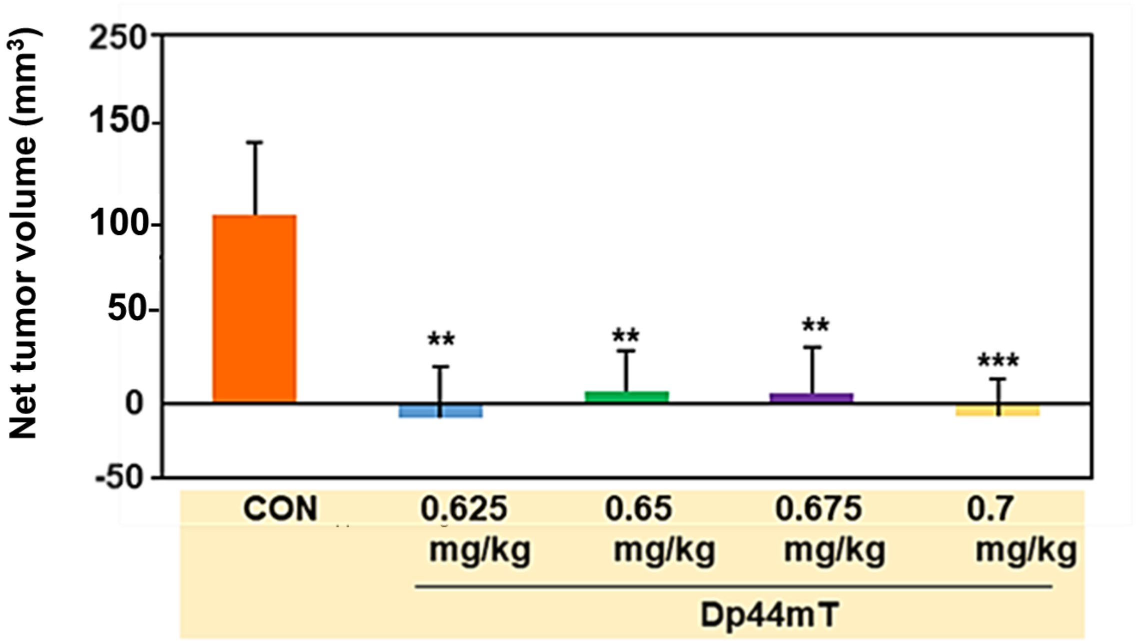


Supplemental Figure 12
